# Mitochondrial remodeling controls innate immune effector function in *Drosophila* hemocytes

**DOI:** 10.64898/2026.03.23.713618

**Authors:** Daewon Lee, Ferdinand Koranteng, Nuri Cha, Sangho Yoon, Kyung-Tae Lee, Jin-Wu Nam, Young V. Kwon, Jiwon Shim

## Abstract

*Drosophila* larval hemocytes, analogous to vertebrate myeloid cells, are central to innate immunity. Despite advances in characterizing hemocyte functions via genetic and biochemical analyses, their metabolic characteristics remain poorly understood. Here, we analyzed metabolic parameters of larval hemocytes across developmental stages, genotypes, and immune activation states using metabolic measurements and single-cell transcriptomics. Our results reveal that hemocytes predominantly rely on mitochondrial respiration for ATP production under homeostasis, with minimal glycolysis. Proliferation or immune activation, particularly lamellocyte differentiation, enhance mitochondrial activity and metabolic flexibility, accompanied by mitochondrial structural remodeling and upregulation of sugar metabolism genes. Functional lamellocytes modulate mitochondrial fission through *Drp1* and utilize glucose and trehalose to fuel mitochondrial respiration, which are essential for effective immune responses. Together, these findings provide a comprehensive profile of hemocyte metabolism and highlight the dynamic interplay between mitochondrial metabolism and innate immunity in invertebrate immune cells.

## Introduction

*Drosophila* blood cells, called hemocytes, are functionally and developmentally analogous to vertebrate myeloid cells^1^. Their conserved yet simple characteristics make hemocytes an attractive invertebrate model for studying hematopoiesis and innate immunity^2^. *Drosophila* hemocytes were initially categorized based on their morphologies^3^ and subsequent studies refined these classifications using marker gene expressions and their biological functions^4^. Most recently, single-cell RNA sequencing (scRNA-seq) analysis allowed a detailed classification of *Drosophila* larval hemocytes, and several studies have provided in-depth insights into the heterogeneity of *Drosophila* hemocyte populations at a transcriptome level^5–9^. Three classic types of mature hemocytes have been identified in *Drosophila* larvae: plasmatocytes, crystal cells, and lamellocytes^2^. Plasmatocytes represent up to 95 % of larval hemocytes and function in phagocytosis, antimicrobial peptide expression, and tissue remodeling, similar to macrophages in vertebrates^10,11^. Crystal cells facilitate wound healing and blood clotting and account for 2 ∼ 5 % of embryonic and larval hemocytes^12,13^. Recently, crystal cells have been shown to regulate internal oxygen homeostasis, in collaboration with the trachea^14^. Lamellocytes are rarely found in healthy animals but arise during immune or stress responses, where they play a role in encapsulation and melanization^15–17^.

Similar to vertebrates, *Drosophila* develops hemocytes by two independent lineages: the embryonic lineage, which originates from the embryonic head mesoderm^10^, and the lymph gland lineage, which arises from the cardiogenic mesoderm^11,18,19^. Embryonically derived hemocytes, known as embryonic hemocytes, proliferate and differentiate primarily into plasmatocytes and migrate throughout the embryo to participate in embryogenesis^10,20,21^. During larval stages, these hemocytes either circulate in the hemolymph or colonize tissue microenvironments, where they continue to proliferate and differentiate^22,23^. The second wave of *Drosophila* hematopoiesis occurs in the lymph gland, which contains hematopoietic progenitors and their microenvironment niche. Prohemocytes in the lymph gland eventually differentiate into mature hemocytes, which are released into circulation following lymph gland disintegration at the onset of pupariation^18,19,24–26^. Hemocytes from both lineages persist through the pupal and adult stages to continue to provide immune surveillance^27^.

An effective immune response demands significant energy, which necessitates enhanced nutritional uptake, metabolic reprogramming of immune cells, and redistribution of body resources. Thus, coordination between metabolism and immunity is critical for effective immune responses and host survival, particularly during infection, across the animal kingdom. In *Drosophila*, the fat body serves as the primary organ for nutrient storage, metabolism, and humoral immunity. Upon immune activation, the fat body allocates stored energy to immune effectors, including hemocytes, to support the acute phase of immune responses^28–30^. It also regulates the number, location, and activation of hemocytes in response to systemic metabolic states^31–33^, thereby controlling overall metabolic homeostasis during immunity. In addition to the fat body, muscles play a key role by modulating whole-animal insulin and JAK/STAT pathways and by storing glycogen required for robust innate immune responses^34–36^. Recent findings have revealed that hemocytes are not merely passive recipients of energy but also act as active regulators, establishing bi-directional communication with other tissues to adjust energy distribution through multiple signaling pathways^30,32,37–39^. This is supported by recent single-cell transcriptome analyses^5–7,9^, which showed that larval hemocyte subtypes express high levels of metabolism-related genes, implying the significance of hemocytes in bridging metabolism and immunity.

Glycolysis and mitochondrial oxidative phosphorylation (OXPHOS) are the two major pathways that generate ATP to support cellular functions. During glycolysis, glucose is converted to pyruvate in the cytosol, producing ATP and reducing equivalents. Pyruvate can subsequently be converted to lactate or transported into mitochondria, where its conversion to acetyl-CoA fuels the tricarboxylic acid cycle (TCA cycle). The TCA cycle generates NADH and FADH2, which donate electrons to the electron transport chain and thereby drive ATP production through OXPHOS. In addition to producing energy, glycolysis and the TCA cycle provide metabolic intermediates required for biosynthesis. Cellular energy metabolism can be assessed in real time using extracellular flux analysis^40^. This technique measures the oxygen consumption rate (OCR) and extracellular acidification rate (ECAR) while serially applying metabolic inhibitors or substrates to modify specific components of cellular metabolism. OCR is commonly used as an indicator of mitochondrial respiration, whereas ECAR reflects extracellular acidification arising from both glycolysis-associated proton release and the hydration of carbon dioxide generated during mitochondrial respiration^41^. Understanding the regulation of metabolic activities provides important insights into how cells adapt their energy metabolism to changing physiological demands.

Despite the growing evidence for metabolic shifts in hemocytes under various developmental or stress conditions, and the significance of hemocytes in the crosstalk between immunity and metabolism, their metabolic profiles under steady-state or immune-activated conditions remain poorly characterized. In this study, we measured metabolic parameters of *Drosophila* larval hemocytes at various developmental timepoints, genotypes, both with and without immune activation. We identified that mitochondrial respiration drives metabolic activation in larval hemocytes. Hemocytes are metabolically dormant under homeostatic conditions but switch to an active state through mitochondrial activation. Furthermore, we demonstrated that lamellocytes exhibit unique metabolic activities, including enhanced trehalose catabolism and mitochondrial remodeling, required for their encapsulation response. Together, our study demonstrates the mitochondria-dependent metabolic activities of *Drosophila* larval hemocytes and their functional relevance in innate immunity.

## Result

### Hemocytes rely on mitochondrial respiration

To understand the metabolic characteristics of embryonically derived hemocytes in larvae, we measured metabolic parameters, including oxygen consumption rate (OCR) and extracellular acidification rate (ECAR), of larval hemocytes using the Seahorse XFe96 Analyzer^42^. For the measurement of hemocytes, we adopted protocols from previous studies^43–45^. First, we successfully reproduced by detecting the previously reported glycolytic signatures in larval brains: Larval brains showed increased ECAR levels following glucose injection, remained unaffected after oligomycin treatment, but significantly decreased upon addition of 2-deoxyglucose (2-DG). These results suggest that glycolysis substantially contributes to extracellular acidification in larval brain cells (Supplementary Figure 1a). Second, based on this protocol, we further optimized the process for hemocyte measurements by changing the culture medium and removing cell debris using a 40 μm cell strainer (see Methods). Lastly, to identify an optimal density of hemocytes in the measurement, we tested different numbers of live hemocytes – 0.43 x 10^6^ cells/mL, 1.13 x 10^6^ cells/mL, and 1.45 x 10^6^ cells/mL – that represent 0.5x, 1x, and 1.5x ratio, and observed proportional increments of OCR or ECAR values in accordance with the absolute number of hemocytes (Figure 1a-b). Based on the manufacturer-recommended basal OCR and ECAR ranges using the Seahorse Xfe96 Analyzer, we set the maximum hemocyte density at 1.5 x 10^6^ cells/mL, corresponding to 2.7 x 10^5^ cells per well in a volume of 180μL. In the absence of drug application, hemocytes maintained constant OCR or ECAR up to 90-minute culture in the Seahorse analyzer, indicating that hemocytes are viable during the metabolic measurements within this time point (Supplementary Figure 1b-c).

**Figure 1.**
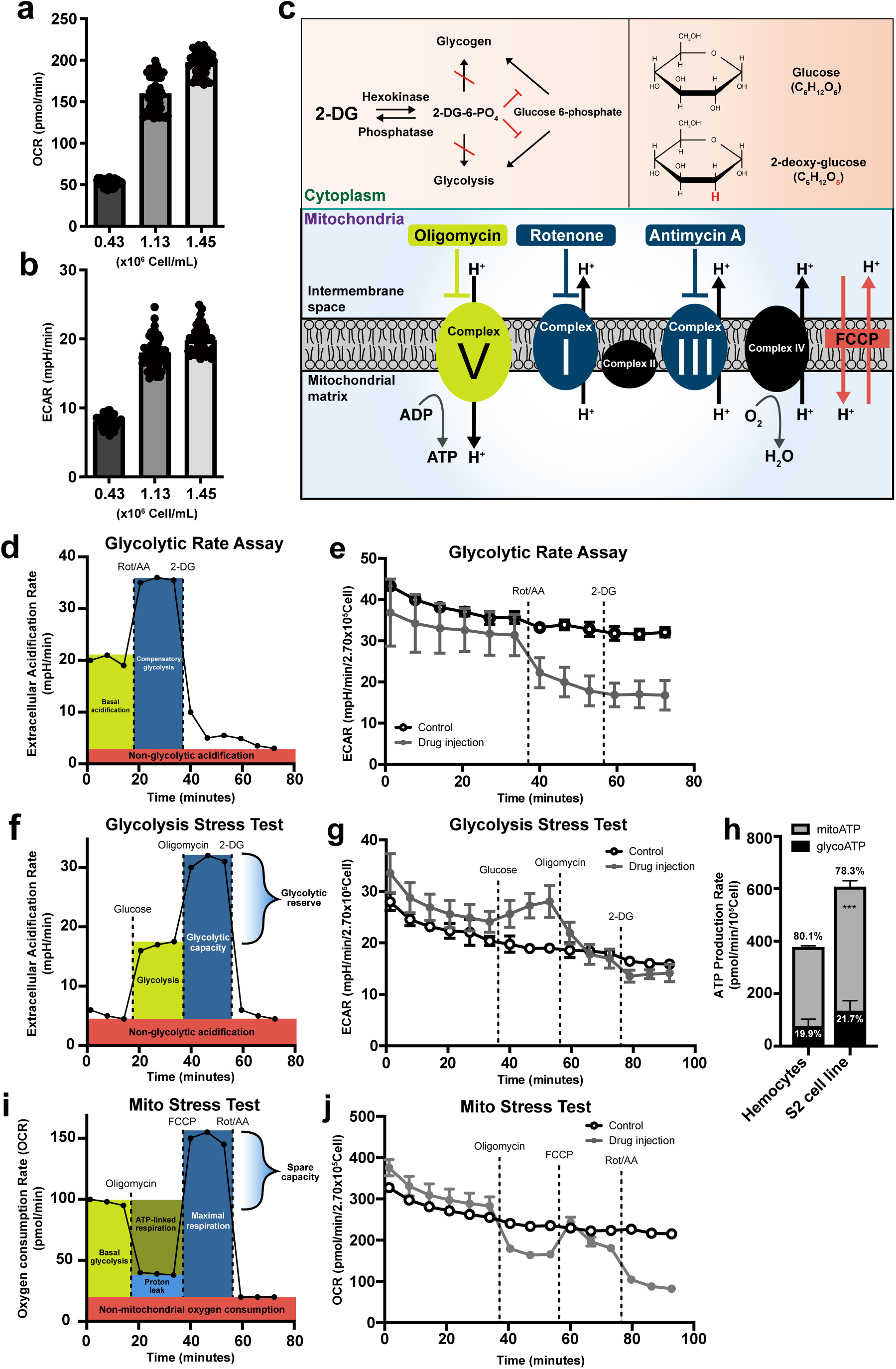
Metabolic analysis of *Drosophila* larval hemocytes. **(a)** Oxygen consumption rates (OCR; pmol/min) and **(b)** extracellular acidification rates (ECAR; mpH/min) of hemocytes (*Hml^Δ^-Gal4, UAS-EGFP*) with different numbers (0.43 x 10^6^ cell/ml, 1.13 x 10^6^ cell/ml, and 1.45 x 10^6^ cell/ml). OCR and ECAR increase in proportion to the number of hemocytes. **(c)** Schematic overview of metabolic inhibitors used in this study. 2-deoxy-glucose (2-DG), a competitive inhibitor of hexokinase, is a glucose analog that lacks the 2-hydroxyl group. Oligomycin inhibits mitochondrial ATP synthase, Complex V. Rotenone or antimycin A inhibit electron transport chain complex I or complex III, respectively. Carbonyl cyanide 4-(trifluoromethoxy)phenylhydrazone (FCCP) is an uncoupling chemical that decreases the proton gradient between mitochondria intermembrane space and mitochondrial matrix. **(d)** Representative Seahorse glycolytic rate assay profile. At baseline, ECAR reflects both glycolytic proton efflux and mitochondrial-derived CO_2_ hydration. Inhibition of mitochondrial respiration with rotenone and antimycin A (Rot/AA) induces ECAR via elevated compensatory glycolysis. Subsequent 2-DG injection suppresses ECAR sustained by glycolysis. The assay medium for Seahorse glycolytic rate assay contains glucose, pyruvate, and glutamine allowing cells to utilize both glycolysis and mitochondrial respiration. **(e)** Seahorse glycolytic rate assay using embryonically derived hemocytes. Rot and AA were injected after the 6^th^ measurement and 2-DG, after the 9^th^ measurement (gray). No glycolytic compensation was observed following Rot/AA treatment. The control group (black) received assay medium at volumes matched to the Rot/AA or 2-DG injections. **(f)** Representative Seahorse glycolysis stress test profile. Glucose was injected to determine glycolysis-induced ECAR levels. Subsequent oligomycin injection inhibits mitochondrial ATP synthase and forces ATP production through glycolysis, indicating glycolytic capacity. Final 2-DG blocks glycolysis. The drop from pre-2-DG ECAR represents the amount of ECAR produced by glycolysis. The remaining ECAR reflects non-glycolytic acidification. The assay medium contains glutamine only (no glucose or pyruvate). **(g)** Seahorse XF glycolysis stress test using embryonically derived hemocytes. Glucose was injected after the 6^th^ measurement, oligomycin after the 9^th^ measurement, and 2-DG after 12^th^ measurement (gray). Glucose injection marginally increased ECAR, which is subsequently blocked by oligomycin treatment. The final 2-DG injection did not change ECAR in hemocytes. The control group (black) received assay medium lacking glucose and pyruvate at volumes matched to the glucose or oligomycin injections. **(h)** ATP production rates of larval hemocytes and S2 cells calculated using the Seahorse real-time ATP rate assay. Hemocytes generated 19.9 % of ATP through glycolysis (glycoATP) and 80.1 % by mitochondrial respiration (mitoATP). S2 cells recapitulated this ratio, producing 21.7 % of glycoATP and 78.3 % via mitoATP. **(i)** Representative Seahorse cell mito stress test profile. The mito stress test measures the key parameters of mitochondrial function, including basal and maximal respiration rates and mitochondrial spare capacities. The oligomycin-induced drop in OCR reflects the ATP-linked mitochondrial respiration and the residual OCR, proton leak. Carbonyl cyanide 4-(trifluoromethoxy)phenylhydrazone (FCCP) injection uncouples the proton gradient across the inner mitochondrial membrane and ATP synthesis that promotes maximal respiration of mitochondria. **(j)** Seahorse XF mito stress test using embryonically derived hemocytes. Oligomycin, FCCP, or Rot/AA was serially injected after the 6^th^, 9^th^, or 12^th^ measurement, respectively (gray). Oligomycin injection reduced OCR, which was recovered by FCCP. Subsequent Rot/AA injection further reduced OCR. Controls (black) were injected with the same volume of mito stress test medium. In **(h)**, statistical analysis was performed using unpaired t-test. n.s: not significant (p > 0.05). *p < 0.05; **p < 0.01; ***p < 0.001. Bars in graphs: the mean. Error bars: standard deviation.

To monitor the mitochondrial activity of hemocytes in the late third instar larvae, we introduced a series of oxidative phosphorylation (OXPHOS) complex inhibitors: oligomycin, rotenone, and antimycin A to hemocytes. Oligomycin, a potent ATP synthase (complex V) inhibitor, impairs the electron flow of the electron transport chain, reducing mitochondrial respiration or OCR^46^. Rotenone or antimycin A is a complex I or complex III inhibitor, respectively, and a mixture of the two compounds shuts down mitochondrial respiration and allows us to measure the non-mitochondrial OCR^47,48^. For measuring glycolysis, 2-deoxy-glucose (2-DG), a glucose analog, was used to competitively block hexokinase and its downstream metabolism^49^ (Figure 1c). First, we carried out the glycolytic rate assay that accounts for compensatory glycolysis following mitochondrial inhibition^50^. Proton efflux in live cells is generated by two main pathways: glycolysis- and mitochondrial-derived acidification^41^. Thus, if tissue is glycolytic, ECAR is significantly elevated after Rot/AA treatment due to compensatory activation of glycolytic proton production (Figure 1d). However, hemocytes considerably reduced ECAR by Rot/AA injection and did not respond to 2-DG (Figure 1e). Also, Rot/AA decreased OCR similar to the trend shown in ECAR (Supplementary Figure 1d), suggesting that OCR and ECAR in larval hemocytes are largely independent of glycolysis. Second, we performed the glycolysis stress test that provides key parameters of glycolytic function, including glycolytic capacity, glycolytic reserve, and non-glycolytic acidification. Most cells are able to switch between glycolysis and oxidative phosphorylation and glucose in cells is converted into pyruvate to be utilized in lactate production in the cytoplasm or TCA cycle. Therefore, glucose injection in a saturated concentration enhances ECAR through lactate dehydrogenase (Ldh)-mediated acidification of cells, which is further elevated by oligomycin treatment due to the lack of OXPHOS^51^ (Figure 1f). Hemocytes marginally induced ECAR upon glucose administration, which was suppressed by oligomycin but not by a following 2-DG injection (Figure 1g). Third, we substantiated these results by real-time ATP rate assay designed to measure total ATP production rates and quantitative bioenergetic balance in cells. Consistent with the glycolytic rate assay and glycolysis test, real-time ATP rate assay demonstrated that larval hemocytes generate ATP mainly by mitochondrial activities (mitoATP, 80.1 %) with a minor contribution of glycolysis (glycoATP, 19.9 %) (Figure 1h; Supplementary Figure 1e-f). In addition to hemocytes from *w^1118^* wild-type animals used for the above experiments, we ran real-time ATP rate assay with hemocyte-derived *Drosophila* S2 cells and hemocytes from different genetic backgrounds, including *Oregon R* and *Hml^Δ^-gal4 UAS-EGFP*, and obtained similar mitochondria-dependent ATP production rates (Figure 1h; Supplementary Figure 1g-k). Finally, mito stress test was used to further investigate a key parameter of mitochondrial functions. Oligomycin decreases electron flow through the electron transport chain (ETC) and FCCP lifts limits of the ETC maximizing oxygen consumption by complex IV (Figure 1i). Mito stress test using hemocytes revealed that FCCP injection did not lead to a rebound in OCR higher than basal levels (Figure 1j), suggesting that hemocytes do not hold the spare mitochondrial capacity, a functional parameter for mitochondrial reserve^52^. Taken together, we conclude that embryonically derived hemocytes in *Drosophila* larvae predominantly utilize mitochondrial respiration to generate energy and exhibit minimal glycolysis rates under unchallenged conditions.

### Metabolic activities of hemocytes change over development

Previous studies have indicated that embryonically derived hemocytes are released into the hemolymph upon hatching and undergo an extensive proliferation during the second to third instars^11,22,53^. Hemocytes actively proliferate during early larval stages, when larvae are actively feeding, whereas their proliferation may decline during the late third instar as larvae approach pupariation. These developmental changes in hemocyte proliferation may be accompanied by corresponding changes in their metabolic activity. We confirmed that the number of embryonically derived circulating hemocytes gradually increases over 72 to 120 h AEL, regardless of dietary sugar composition (Supplementary Figure 2a). Based on this developmental profile, we next examined whether hemocytes undergo corresponding metabolic fluctuations throughout larval development. To assess developmental changes in larval hemocyte metabolism during the second and third instars, we measured OCR or ECAR of larval hemocytes at 72 h, 96 h, or 120 h after egg laying (AEL), representing mid-second, early third, and late third instars, respectively. With the real-time ATP rate assay, we observed that hemocytes at 96 h AEL generate larger ATP production rate than those in 72 h or 120 h AEL (Figure 2a, Supplementary Figure 2b-c). In addition, we examined an intermediate developmental time point at 108 h AEL and found that the number of circulating hemocytes increased from 96 to 120 h AEL, including 108 h AEL (Supplementary Figure 2d), whereas mitochondrial activity and ATP production rates gradually decreased over the same developmental period (Supplementary Figure 2e-g). At all stages, ATP production relied heavily on mitochondrial respiration, and the proportion of glycolysis-mediated ATP generation remained low, which rather diminished over development (Figure 2a; Supplementary Figure 2g). Consistently, ATP-linked respiration corresponded well with the ATP production rate (Figure 2b). This phenotype was recapitulated by biochemical ATP measurement assay (Supplementary Figure 2c). Proton leak, an indicator for mitochondrial fitness, decreased over time (Figure 2c). Furthermore, hemocytes did not respond to 2-DG at all developmental time points (Supplementary Figure 2h). These results indicate that mitochondrial respiration is the main process for hemocyte metabolism and ATP production peaks at 96 h AEL during hemocyte development.

**Figure 2.**
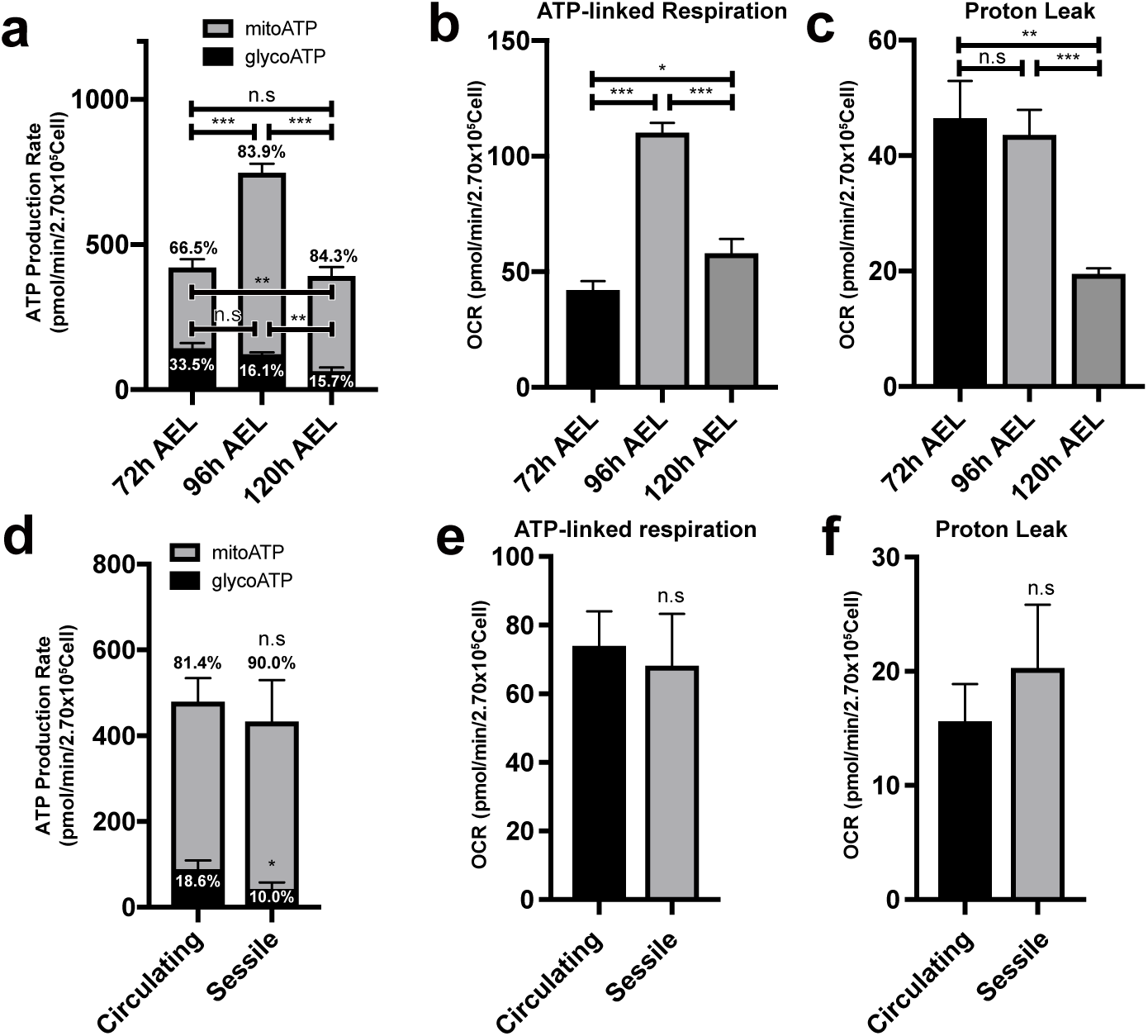
Metabolic variations of hemocytes during larval development. **(a)** ATP production rate of *Drosophila* larval hemocytes at different developmental timepoints. Hemocytes at 72 h after egg laying (AEL) produce 33.5% ATP by glycolysis and 66.5 % by mitochondrial respiration. At 96 h AEL, hemocytes exhibit 16.1 % of glycoATP and 83.9% of mitoATP production rates and generate larger amount of ATP than 72 h or 120 h. Hemocytes at 120 h AEL express 15.7 % of glycoATP and 84.3 % of mitoATP production rates. The ratio of glycoATP reduces over development. **(b)** ATP-linked respiration levels of hemocytes at 72 h, 96 h, and 120 h AEL. OCR is the highest at 96 h AEL. **(c)** Proton leak levels of larval hemocytes at different developmental timepoints. Proton leak gradually decreases during development. **(d)** ATP production rates of circulating or sessile hemocytes. Circulating hemocytes display 18.6 % of glycoATP and 81.4 % of mito ATP production rates. Sessile hemocytes show 10.0 % of glycoATP and 90.0 % of mitoATP production rates. **(e)** ATP-linked respiration levels of circulating or sessile hemocytes. ATP-linked respiration levels are indistinguishable in circulating or sessile hemocytes. **(f)** Proton leak levels of circulating or sessile hemocytes. Circulating or sessile hemocytes exhibit similar proton leak levels. In **(a)** to **(f)** statistical analysis was performed using unpaired t-test. n.s: not significant (p > 0.05). *p < 0.05; **p < 0.01; ***p < 0.001. Bars in graphs: the mean. Error bars: standard deviation.

Upon hatching, larval hemocytes either freely circulate the larval hemolymph or colonize resident tissues including hematopoietic pockets of the larval body wall to control their proliferation and differentiation^54^. Considering that hemocyte metabolism is modified during larval development, we asked whether circulating or sessile hemocytes display differential cellular metabolism due to their distinct microenvironment. Glycolysis-mediated ATP production was lower in sessile hemocytes (10 %) than in circulating hemocytes (18.6 %) (Figure 2d). However, most metabolic profiles, such as ATP production, ATP-linked respiration, and proton leak, were comparable regardless of the hemocyte location (Figure 2e-f; Supplementary Figure 2i-j). Overall, we conclude that hemocytes developmentally elevate ATP production rates at 96 h AEL and that hemocytes at two locations – sessile and circulating – are metabolically indistinguishable.

### Hemocyte proliferation and differentiation are associated with mitochondrial respiration

Our findings indicate that larval hemocytes rely on mitochondrial respiration without reserving spare mitochondrial capacities. However, metabolic activities of hemocytes fluctuate during larval development, implying that these cells are able to transform cellular metabolism according to bioenergetic requirements. Based on this, we sought to understand the metabolic signatures of hemocytes in various genetic backgrounds that promote the proliferation or differentiation of hemocytes or upon immune challenges. First, we investigated whether induced proliferation or activation of the MAPK pathway alters hemocyte metabolism by using an active form of *Ras*, *UAS-Ras^v12^* (*Hml^Δ^-Gal4 UAS-Ras^v12^*). We confirmed that late-third instar larvae carrying both transgenes show over a 10-fold increase in the number of *Hml^+^* plasmatocytes with a relatively minor proportion of lamellocytes differentiation (Supplementary Figure 3a-d). Despite the stark increase in hemocyte numbers, *Ras^v12^* expression caused only a modest increase in the proportion of pH3-positive hemocytes at the late third instars (Supplementary Figure 3e). A fixed number of hemocytes (2.7 x 10^5^ cells) expressing *Ras^v12^* exhibited 2-fold higher basal OCR or ECAR levels than those in controls, which was significantly decreased to non-mitochondrial OCR levels following oligomycin and Rot/AA injections (Supplementary Figure 3f-g). Additionally, the expression of *Ras^v12^* considerably enhanced the ATP production rate without changing the preference for mitochondrial respiration over glycolysis (Figure 3a). Both ATP-linked respiration (Figure 3b) and proton leak (Figure 3c) were increased in this genetic background. Different from wild-type hemocytes, hemocytes expressing *Ras^v12^* showed higher OCR rebound than basal levels following the FCCP treatment (Figure 3d), indicating that *Ras^v12^*-expressing hemocytes elevate the spare respiratory capacity. Thus, we conclude that *Ras^v12^*-expressing hemocytes rewire mitochondrial capacities as well as energetic activities.

**Figure 3.**
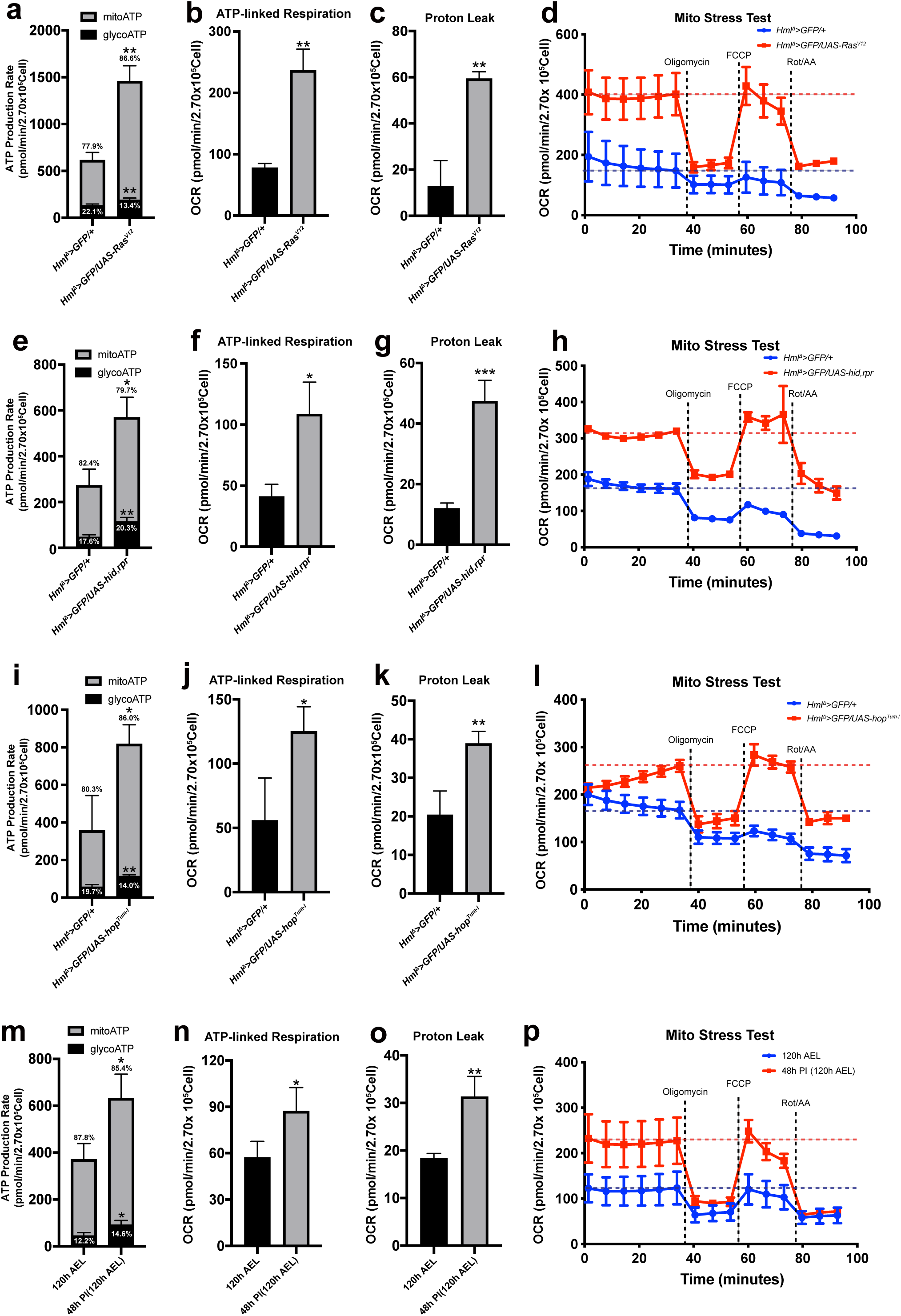
Metabolic changes of hemocytes in multiple genetic backgrounds or upon wasp infestation. **(a)** ATP Production Rate of wild-type (*Hml^Δ^-Gal4, UAS-EGFP/+*) or *Ras^v12^*-expressing hemocytes (*Hml^Δ^-Gal4, UAS-EGFP; UAS-Ras^v12^*) calculated by Seahorse XF real-time ATP rate assay. Wild-type hemocytes display 22.1 % of glycoATP and 77.9 % of mito ATP production rates while hemocytes expressing *Ras^v12^* exhibit 13.4 % of glycoATP and 86.6 % of mitoATP. *Ras^v12^*-expressing hemocytes express higher ATP production rates than control. **(b)** ATP-linked respiration levels of wild-type hemocytes (*Hml^Δ^-Gal4, UAS-EGFP/+*) or *Ras^v12^*-expressing hemocytes (*Hml^Δ^-Gal4, UAS-EGFP; UAS-Ras^V12^*). **(c)** Proton leak levels of hemocytes of control (*Hml^Δ^-Gal4, UAS-EGFP/+*) or *Ras^v12^*-expressing hemocytes (*Hml^Δ^-Gal4, UAS-EGFP; UAS-Ras^V12^*). **(d)** Mito stress test measuring OCR levels of control (*HmlΔ-Gal4, UAS-EGFP/+*; blue) or *Ras^v12^*-expressing hemocytes (*Hml^Δ^-Gal4, UAS-EGFP;UAS-Ras^V12^*; red). **(e)** ATP production rate of control hemocytes (*Hml^Δ^-Gal4, UAS-EGFP/+*) or hemocytes remaining after ablation of the Hml^+^ population (*Hml^Δ^-Gal4, UAS-EGFP; UAS-hid,rpr*) calculated by Seahorse XF real-time ATP rate assay. Control hemocytes express 17.6% of glycoATP and 82.4% of mito ATP production rates. Genetic ablation of *Hml^+^*hemocytes leads to 20.3% of glycoATP and 79.7% of mitoATP production rates with higher ATP production rates than control. **(f)** ATP-linked respiration levels in control hemocytes (*Hml^Δ^-Gal4, UAS-EGFP/+*) and hemocytes remaining after ablation of the Hml^+^ population (*Hml^Δ^-Gal4, UAS-EGFP; UAS-hid,rpr*). **(g)** Proton leak levels in control hemocytes (*Hml^Δ^-Gal4, UAS-EGFP/+*) and hemocytes remaining after ablation of the Hml^+^ population (*Hml^Δ^-Gal4, UAS-EGFP; UAS-hid,rpr*). **(h)** Mito stress test measuring OCR levels in control hemocytes (*Hml^Δ^-Gal4, UAS-EGFP/+*; blue) and hemocytes remaining after ablation of the Hml^+^ population (*Hml^Δ^-Gal4, UAS-EGFP; UAS-hid,rpr*; red). **(i)** ATP production rate of control (*Hml^Δ^-Gal4, UAS-EGFP/+*) or hemocytes expressing *hop^Tum-l^* (*Hml^Δ^-Gal4, UAS-EGFP; UAS-hop^Tum-l^*) measured by Seahorse XF real-time ATP rate assay. Control hemocytes express 19.7 % of glycoATP and 80.3 % of mito ATP production rates. Overexpression of *hop^Tum-l^* in hemocytes results in 14.0 % of glycoATP and 86.0 % of mitoATP production rate and higher ATP production rates. **(j)** ATP-linked respiration of Control (*Hml^Δ^-Gal4, UAS-EGFP/+*) or *hop^Tum-l^*-expressing hemocytes (*Hml^Δ^-Gal4, UAS-EGFP; UAS-hop^Tum-l^*). **(k)** Proton leak levels of control (*Hml^Δ^-Gal4, UAS-EGFP/+*) or *hop^Tum-l^*-expressing hemocytes (*Hml^Δ^-Gal4, UAS-EGFP; UAS-hop^Tum-l^*). **(l)** Mito stress test measuring OCR levels of control (*Hml^Δ^-Gal4, UAS-EGFP/+*; blue) or *hop^Tum-l^* -expressing hemocytes (*Hml^Δ^-Gal4, UAS-EGFP; UAS-hop^Tum-l^*; red). **(m)** ATP production rate of hemocytes (*Hml^Δ^-Gal4, UAS-EGFP*) at 120 h AEL or 48 h post-infestation (48 h PI; equivalent to 120 h AEL) measured by Seahorse XF real-time ATP rate assay. Wild-type hemocytes express 12.2 % of glycoATP and 87.8 % of mito ATP production rates. Immune-activated hemocytes show 14.6 % of glycoATP and 85.4 % of mitoATP production rates with higher ATP production rates than unchallenged controls. **(n)** ATP-linked respiration levels of control hemocytes at 120 h AEL or 48 h PI. **(o)** Proton leak levels of hemocytes at 120 h AEL or 48 h PI. **(p)** Mito stress test measuring OCR levels of hemocytes at 120 h AEL (blue) and 48 h PI (equivalent to 120 h AEL; red). In **(d)**, **(h)**, **(l)**, and **(p)**, oligomycin, FCCP, and Rot/AA were injected after 6^th^, 9^th^, or 12^th^ measurement, respectively. Blue or red dashed line indicates average OCR levels from 1^st^ to 6^th^ cycle measurement. In **(a)-(c)**, **(e)-(g)**, **(i)-(k),** and **(m)-(o)**, statistical analysis was performed using unpaired t-test. n.s: not significant (p > 0.05). *p < 0.05; **p < 0.01; ***p < 0.001. Bars in graphs: the mean. Error bars: standard deviation.

Recent studies including single-cell transcriptome analyses of larval hemocytes indicated that larval plasmatocytes are composed of transcriptionally diverse subtypes, in addition to the *Hml^+^* population^5,6^. To test whether every plasmatocyte, regardless of its subtype represented by a marker gene expression, displays similar metabolic profiles during proliferation, we expanded *Hml^-^*plasmatocytes by genetically ablating *Hml^+^* plasmatocytes. Expression of the pro-apoptotic genes *hid* and *rpr* in *Hml^+^* plasmatocytes (*Hml^Δ^-Gal4 UAS-hid,rpr*) reduces the number of *Hml^+^* plasmatocytes while expanding *Hml^-^*but *Pxn^+^* plasmatocytes^55^. Similar to *Ras^v12^*-expressing *Hml^+^* plasmatocytes, *Hml^-^Pxn^+^* plasmatocytes exhibited higher basal OCR or ECAR rates (Supplementary Figure 3h-i). These rates sharply declined following oligomycin and Rot/AA treatment in a similar trend seen by those expressing *Ras^v12^* (Supplementary Figure 3f-g). Consistently, ATP production rates (Figure 3e), ATP-linked respiration (Figure 3f), proton leak (Figure 3g), and the spare respiratory capacity (Figure 3h) were all increased. These results suggest that proliferation of plasmatocytes, regardless of their subtypes, augments hemocyte metabolism. It is worth noting that the expansion of specific hemocyte subtypes may be associated with cell death-induced inflammation and could therefore be biologically distinct from *Ras^v12^*-mediated hemocyte proliferation

Lastly, we assessed the metabolic activities of crystal cells, the second largest population under normal growing conditions. Since crystal cells comprise only ∼ 5 % of total hemocytes, we increased their proportion by accelerating crystal cell differentiation through *Notch^ICD^* expression (*Hml^Δ^-Gal4 UAS-N^ICD^*). Overexpression of *Notch^ICD^* successfully expanded the crystal cell pool (Supplementary Figure 3j). However, number-matched *Notch^ICD^*-expressing hemocytes displayed marginally decreased or similar metabolic levels comparable to control hemocytes, including ATP production, ATP-linked respiration, and proton leak (Supplementary Figure 3k-m). Moreover, the mito stress test was unchanged in this genetic background (Supplementary Figure 3n), suggesting *N^ICD^*-expressing hemocytes with expanded crystal cell population exhibit lower metabolic activities than wild-type hemocytes. Overall, these results establish that an increase in the number of plasmatocytes, but not crystal cells, is accompanied by enhanced respiratory activity and increased mitochondrial reserve.

### Lamellocyte differentiation or active immunity triggers mitochondrial respiration

While plasmatocytes are the major hemocyte population under unchallenged conventional conditions, lamellocytes are the predominant cell type that give rise during active immunity or stress responses^15,16^. To gain further insights into the lamellocyte differentiation and mitochondrial function, we expressed a constitutively active form of the Janus kinase Hopscotch (*hop^Tum-l^*) in hemocytes (*Hml^Δ^-gal4 UAS-hop^Tum-l^*) to genetically induce a large number of lamellocytes (Supplementary Figure 4a-c)^17,56,57^. Hemocyte-specific expression of *hop^Tum-l^* considerably elevated basal OCR or ECAR and raised ATP production rates, ATP-linked respiration, and proton leak (Figure 3i-k; Supplementary Figure 4d-e). Also, we detected spare respiratory capacities in *hop^Tum-l^* expressing hemocytes similar to those found in *Ras^v12^*- or *hid, rpr*-expressing hemocytes (Figure 3l).

*Drosophila* larvae respond to wasp-egg deposition within a few hours and trigger a series of innate immune responses, including a massive differentiation of lamellocytes^58^. Immature lamellocytes appear by 24 h post-infestation (PI) and develop into mature lamellocytes by 48 h PI to encapsulate wasp eggs^5,59^. Given that proliferation or differentiation of hemocytes stimulated by gene expression alters hemocyte metabolism, we asked whether innate immunity caused by a natural infection influences hemocyte respiration. Hemocytes at 24 h PI when mature lamellocytes have not yet given rise showed OCR or ECAR levels similar to unchallenged controls at 96 h AEL and did not change other metabolic indicators (Supplementary Figure 4f-g). On the other hand, hemocytes at 48 h PI expressed significantly higher OCR or ECAR by two folds than wild-type hemocytes at 120 h AEL (Supplementary Figure 4h-i). Moreover, these hemocytes increased their ATP production rate, ATP-linked mitochondrial respiration, and proton leak (Figure 3m-o). The mito stress test recognized that hemocytes at 48 h PI result in elevated spare respiratory capacities (Figure 3p), suggesting that immune-activated hemocytes and mature lamellocytes amplify mitochondrial respiratory rates and capacity.

Overall, these data suggest that while larval hemocytes under unchallenged conditions rely largely on mitochondrial respiration and exhibit low metabolic activity, hemocyte proliferation, lamellocyte differentiation, or active immune responses significantly elevate mitochondrial as well as glycolytic activities to promote ATP production.

#### Mitochondrial activation promotes hemocyte proliferation and lamellocyte differentiation

Based on our observations that genetically induced proliferation and differentiation are accompanied by increased mitochondrial activity and ATP production, we asked whether enhanced mitochondrial activity itself is sufficient to promote hemocyte proliferation and differentiation. TCA cycle flux is partly influenced by the availability of acetyl-CoA, which is generated from pyruvate by the pyruvate dehydrogenase complex (PDH)^60^. PDH activity is negatively regulated by pyruvate dehydrogenase kinase (*Pdk*), which phosphorylates and inactivates PDH. Therefore, *Pdk* RNAi is expected to increase acetyl-CoA production and TCA cycle activity, potentially enhancing mitochondrial electron transport chain^61^. Expression of *Pdk* RNAi in hemocytes (*Srp-Gal4 Hml^Δ^-Gal4 UAS-Pdk RNAi*) increased circulating hemocyte numbers and promoted lamellocyte differentiation during normal development (Figure 4a-b).

**Figure 4.**
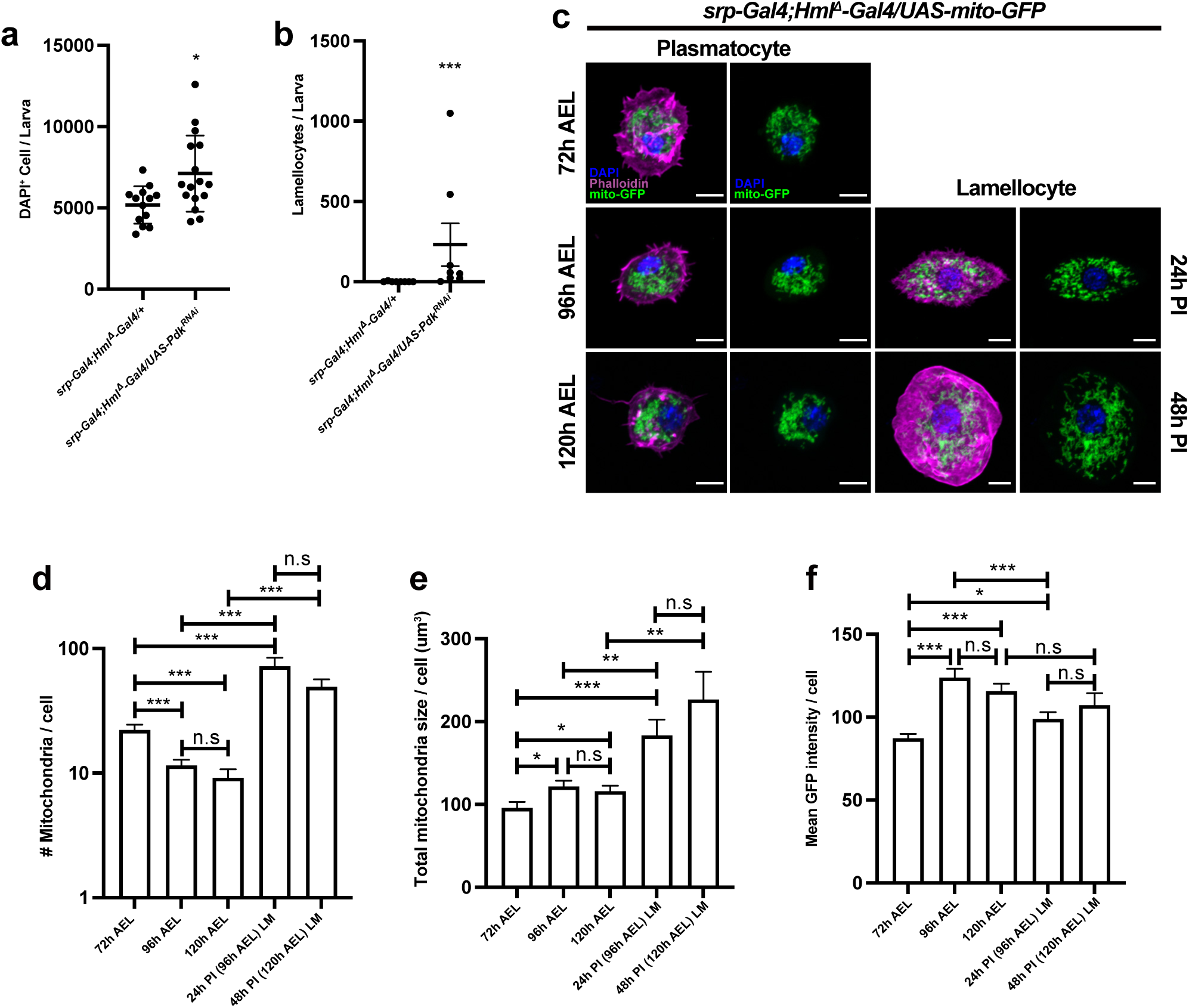
Pdk knockdown and mitochondrial remodeling in lamellocyte differentiation. **(a)** Quantification of DAPI-positive embryonic hemocytes in *srp-Gal4;Hml^Δ^-Gal4/+* and *srp-Gal4;Hml^Δ^-Gal4/UAS-Pdk RNAi*. Bars in graphs: the mean. Error bars: the standard deviation. **(b)** Quantification of lamellocytes among embryonic hemocytes in *srp-Gal4;Hml^Δ^-Gal4/+* and *srp-Gal4;Hml^Δ^-Gal4/UAS-Pdk RNAi*. Bars in graphs: the mean. Error bars: the standard error of the mean. **(c)** Mitochondria morphology of plasmatocytes or lamellocytes at different developmental time points and upon wasp infestation (*srp-Gal4;Hml^Δ^-Gal4/UAS-mito-GFP*). Larvae were infected by wasps at 72 h AEL; thus, no corresponding lamellocyte image at this timepoint. 96 h AEL corresponds to 24 h post-infection (PI). 120 h AEL, 48 h PI. Phalloidin, magenta; mito-GFP; green, DAPI, blue. Scale bar: 5 μm (white). **(d)** The number, **(e)** size of mitochondria per cell, and **(f)** mito-GFP intensity per cell at different developmental time points or upon wasp infection. Bars in graphs: the mean. Error bars: the standard error of the mean. In (**a)**, (**b)**, (**d), (e)** and (**f),** statistical analysis was performed using unpaired t-test. n.s: not significant (p > 0.05). *p < 0.05; **p < 0.01; ***p < 0.001.

Consistent with the enhanced mitochondrial metabolic activities following wasp parasitism (Figure 3m-p), we found that tetramethylrhodamine (TMRM), an indicator of mitochondrial membrane potential, and dihydroethidium (DHE), a superoxide sensor, were both marginally increased in hemocytes (Supplementary Figure 5a-b). These findings suggest that wasp infestation is accompanied by increased mitochondrial membrane potential and intracellular ROS levels, consistent with changes in mitochondrial activities during immune activation. To determine whether elevated ROS levels alone are sufficient to alter hemocyte development, we manipulated intracellular ROS levels using genetic perturbations. Increasing intracellular ROS levels through hemocyte-specific knockdown of *ND75*, a core subunit of the mitochondrial complex I (*Hml^Δ^-Gal4 UAS-ND75 RNAi*)^62^ did not significantly alter the number of GFP-positive circulating hemocyte but marginally increased the total numbers of hemocytes and lamellocytes (Supplementary Figure 5c-e). Furthermore, succinate supplementation did not affect the total number of circulating hemocytes, although it promoted lamellocyte differentiation even in the absence of wasp infection (Supplementary Figure 5f-h). Overall, these results indicate that enhancing PDH-dependent mitochondrial metabolism promotes both circulating hemocyte accumulation and lamellocyte differentiation. Together with our observation that genetically induced hemocyte proliferation and differentiation are accompanied by increased mitochondrial respiration, these findings suggest a close and potentially bidirectional relationship between hemocyte development and mitochondrial metabolic activity.

### Rearrangement of mitochondrial structures in plasmatocytes and lamellocytes

Since metabolic alterations often accompany structural modifications of mitochondria^63^, we hypothesized that mitochondria in hemocytes undergo morphological or quantitative changes during hemocyte development or upon active immunity. To address this, we used *UAS-mitoGFP* to visualize mitochondrial morphology in embryonically derived hemocytes (*srp-Gal4;Hml^Δ^-Gal4/UAS-mitoGFP*). Over larval development under unchallenged conditions, plasmatocytes experienced a significant increase in mitochondrial volume during 72 to 96 h AEL and maintained the volume until the third instar 120 h AEL (Figure 4c-f). In contrast, the number of mitochondria per one plasmatocyte decreased, particularly between 72 and 96 h AEL (Figure 4c-f). A stark volumetric change during 72 to 96 h AEL corresponded well with fluctuations of metabolic parameters of hemocytes at this stage (Figure 2a-c), supporting that metabolic plasticity is associated with mitochondrial rearrangement. Under wasp parasitism, immature lamellocytes given rise at 24 h PI contained a higher number of mitochondria with a significantly smaller size than age-matched plasmatocytes at 96 h AEL (Figure 4c-f). At 48 h PI, mature lamellocytes marginally reduced the number of mitochondria and keep the mitochondrial volume comparable to immature lamellocytes at 24 h PI (Figure 4c-f). These results suggest two different regulatory modes of mitochondrial plasticity in plasmatocytes and lamellocytes: developing plasmatocytes maintain relatively constant numbers of mitochondria with an increase in size during the second to third instar. However, lamellocytes undergo extensive mitochondrial fragmentation during the initial differentiation process, which is later attenuated and shifted towards the size expansion during maturation.

### Mitochondrial fission is required for lamellocyte immune function

To understand how hemocytes acquire differential mitochondrial activity during development and immune activation, we examined transcriptional profiles of metabolic genes in wild-type and wasp-infested larval hemocytes extracted from the single-cell RNA sequencing^5,7^. We integrated single-cell RNA transcriptomes of hemocytes into pseudo-bulk levels and clustered cell types into five groups: developing plasmatocytes found in conventional conditions (PM), plasmatocytes upon wasp infestation (PM_inf), developing crystal cells in conventional conditions (CC), crystal cells upon wasp infestation (CC_inf), and lamellocytes (LM_inf). Plasmatocytes changed their metabolic gene expressions during wasp parasitism by downregulating *Hex*-*A*, *UQCR-6.4* (mitochondrial electron transport, complex III), *ND-AGGG* (mitochondrial electron transport, complex I), and up-regulating *Gapdh2* or *ATPsynbeta* (Figure 5a). Moreover, crystal cells moderately modified similar gene sets upon wasp infestation. Lamellocytes, on the other hand, displayed considerably distinct metabolic gene expression profiles compared with plasmatocytes. Four representative categories of genes were highly induced in lamellocytes: glucose homeostasis (*Gapdh1*, *Gapdh2*, *Tpi*, *Pglym78*, and *Eno*), mitochondrial fission (*Drp1* and *Fis1*), SLC2 family sugar transporter (*CG1208*, *Tret1-1*, and *CG4607*), and starch and sucrose metabolism (*Treh*) (Figure 5a). Consistent with the scRNA-seq analysis, RT-qPCR analysis of hemocyte mRNA supported increased expression of representative metabolic genes upon wasp infestation (Supplementary Figure 8a-k). Notably, mitochondrial fission genes, including *Drp1* and *Fis1* were progressively upregulated in lamellocytes from 24h PI, compared with corresponding their expression in plasmatocytes at 96h AEL, reaching peak expression at 48h PI (Supplementary Figure 6a). This temporal expression pattern coincided with changes in mitochondrial morphologies in differentiated lamellocytes (Figure 4c-f). However, we found no apparent changes in the mitochondrial fusion gene, *Marf*, in lamellocytes (Supplementary Figure 6b). We confirmed that *Drp1* knockdown in lamellocytes at 48 h PI significantly impaired mitochondrial fission and resulted in highly elongated mitochondria (*Srp-Gal4 UAS-mitoGFP UAS-Drp1 RNAi*) (Supplementary Figure 6c). Moreover, hemocytes with reduced *Drp1* expression decreased mitochondrial activities compared with controls (Supplementary Figure 6d-e). To test whether mitochondrial fission is associated with lamellocyte differentiation or function, we analyzed lamellocyte numbers and encapsulation success rates after knocking-down *Drp1* in hemocytes (*Hml^Δ^-Gal4 UAS-GFP UAS-Drp1 RNAi*). Despite the marked downregulation of *Drp1* mRNA (Supplementary Figure 6f), *Drp1* RNAi did not alter lamellocyte differentiation (Supplementary Figure 6g-h). However, hemocyte-specific expression of *Drp1* RNAi significantly reduced encapsulation of wasp eggs (Figure 5b-c; Supplementary Figure 6i), suggesting that mitochondrial remodeling is required for melanization response mediated by lamellocytes. Together, these findings indicate that relative to both unchallenged and challenged plasmatocytes, lamellocytes emerging upon wasp infestation exhibit differential expression of genes associated with sugar metabolism and mitochondrial structure and activity. Additionally, morphological rearrangement of mitochondria is critical for immune effector function of lamellocytes (Figure 5d).

**Figure 5.**
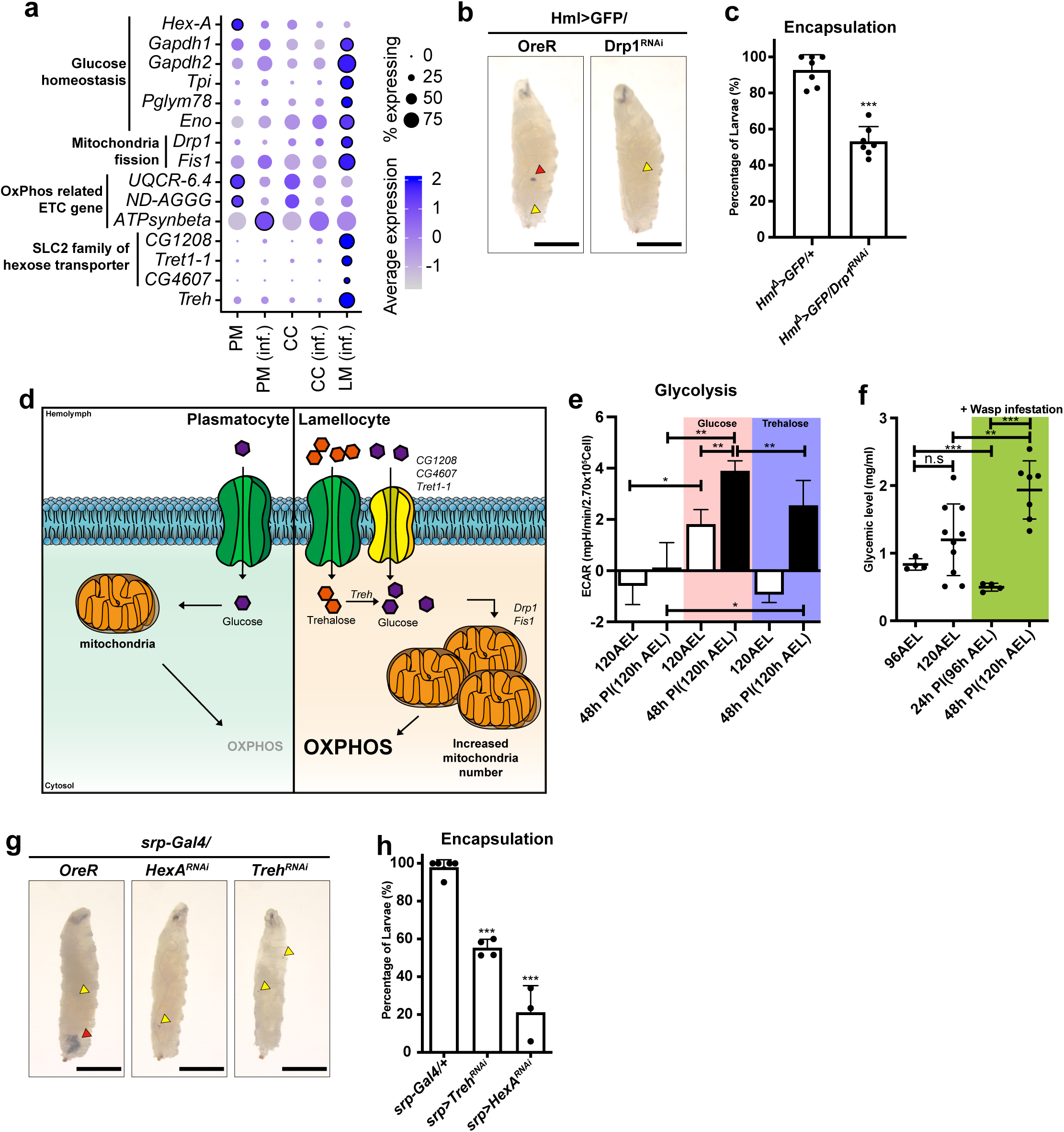
Mitochondrial modification in hemocytes upon wasp infestation. **(a)** Dot plot expression of significantly changed metabolic genes across cell clusters. PM: plasmatocytes under conventional conditions, PM (inf.): plasmatocytes under wasp infestation, CC: crystal cells under conventional conditions, CC (inf.): crystal cells under wasp infestation, LM (inf.): lamellocytes under wasp infestation. Dot size represents the percentage of cells expressing each gene. Dot color shows average expression levels of metabolic genes. **(b)** Encapsulation of wasp eggs in *Drosophila* larvae at 60 h post-infestation (60 h PI). Control larvae (*Hml^Δ^-Gal4, UAS-EGFP/+*) containing melanized wasp eggs (red arrowhead) (left). Hemocyte-specific *Drp1* RNAi (*Hml^Δ^-Gal4, UAS-EGFP, Drp1 RNAi*) reduces the rate of wasp egg encapsulation (right). Yellow arrowhead marks the wasp ovipositor injection site. Scale bar: 1 mm (black) **(c)** Quantification of encapsulation rates shown in **(b)**. Each dot represents the encapsulation ratio from each trial (n=147, 230 respectively). **(d)** A schematic diagram correlating metabolic gene expressions, mitochondrial morphology, and metabolic modification in plasmatocytes and lamellocytes. After wasp infestation hemocytes differentiate into lamellocytes and increase the expression of sugar transporters (*CG1208, CG4607 and Tret1-1*) and trehalase (*Treh*) to enhance sugar uptake. In addition, lamellocytes increase both the number and size of mitochondria, leading to an increased capacity for oxidative phosphorylation (OXPHOS). **(e)** Quantification of ECAR following glucose or trehalose injection corresponding to supplementary figure 6(j) and **(k).** Pink shade indicates glucose-injected samples; purple indicates trehalose. Black bar denotes hemocytes upon 48 h post-infection (corresponding to 120 h AEL). **(f)** Glycemic levels of larval hemolymph at different developmental time points (96 h AEL and 120 h AEL) and infestation condition (24 h PI and 48 h PI). Green shading indicates wasp infested conditions. **(g)** Encapsulation of wasp eggs in *Drosophila* larvae at 60 h post-infestation (60 h PI). Wild-type larvae containing melanized wasp eggs (red arrowhead) (left). Hemocyte-specific *Hex-A* RNAi (*srp-Gal4 UAS-Hex-A RNAi*) or *Treh* RNAi (*srp-Gal4 UAS-Treh RNAi*) reduced the rate of wasp egg encapsulation (middle and right). Yellow arrowhead marks the wasp ovipositor injection site. Scale bar: 1 mm (black) **(h)** Quantification of encapsulation rates shown in **(g)**. Each dot represents the encapsulation ratio from 40 animals (n=200, 160, 120, respectively). In (**c), (e)**, **(f)** and (**h),** statistical analysis was performed using unpaired t-test. n.s: not significant (p > 0.05). *p < 0.05; **p < 0.01; ***p < 0.001. Bars in graphs: the mean. Error bars: standard deviation.

### Functional lamellocytes utilize sugars to fuel mitochondrial respiration

Given that hemocytes at 48 h PI stimulate energy production and lamellocytes express increased levels of sugar metabolism and mitochondria-related genes, we investigated whether hemocytes at 48 h PI respond differently to sugars. Trehalose, a primary sugar type used by *Drosophila*, is either stored in the fat body or mobilized to glucose as an energy source^64^. To test if hemocytes under unchallenged conditions and immune-activated hemocytes differentially respond to sugars, we injected trehalose or glucose into control hemocytes at 120 h AEL or lamellocytes at 48 h PI. Hemocytes at 120 h AEL marginally elevated ECAR levels after glucose inoculation, which subsequently declined following oligomycin administration (Figure 5e; Supplementary Figure 6j). However, trehalose injection did not raise ECAR levels in control hemocytes, implying a specific response of developing hemocytes to glucose (Figure 5e; Supplementary Figure 6j). In contrast, hemocytes at 48 h PI, mainly comprised of lamellocytes, reacted to both glucose and trehalose injection by instantly increasing their ECAR levels, which were suppressed by oligomycin (Figure 5e; Supplementary Figure 6k). The significance of glycolysis in the ECAR elevation was probed by applying 2-DG to these hemocytes following the oligomycin treatment. 2-DG did not alter ECAR similarly to unchallenged controls (Supplementary Figure 6j-k). These data suggest that lamellocytes utilize glucose and trehalose mainly through mitochondrial respiration rather than activating glycolysis.

Innate immunity counteracts nutrient storage^65,66^ and consequently changes the hemolymph sugar levels^39,67,68^. To understand how systemic sugar levels correlate with hemocyte metabolism, we measured hemolymph sugar levels before and after wasp infestation and across development. Between 96 to 120 h AEL, larvae showed a dramatic increase in glycemic levels that represent both trehalose and glucose in the hemolymph (Figure 5f). Similar to developing hemocytes, wasp infestation led to a robust increase in glycemic levels at 48 h PI, reaching levels higher than those observed in 120h AEL (Figure 5f). To test whether elevated glycemia and sugar responsiveness during wasp parasitization contribute to hemocyte differentiation or proliferation, we quantified total hemocytes, Pxn^+^ plasmatocytes, and lamellocytes after hemocyte-specific RNAi against *HexA* or *Treh* RNAi (*Srp-Gal4 UAS-Treh RNAi* or *Srp-Gal4 UAS-HexA RNAi*). Knockdown of these sugar metabolism or transporter genes did not alter absolute hemocyte counts, including lamellocytes (Supplementary Figure 7a-c). However, it significantly reduced the efficiency of wasp egg encapsulation (Figure 5g-h), indicating that sugar metabolism is essential for the functionality of lamellocyte-mediated encapsulation, similar to the inhibition of mitochondrial fission (Figure 5b-c). Together, these results indicate that developing hemocytes are selectively responsive to glucose, whereas hemocytes stimulated during wasp parasitism (48h PI), which largely comprise lamellocytes, utilize both glucose and trehalose to support elevated metabolic activities. Furthermore, metabolic reprogramming, associated with mitochondrial remodeling and sugar responsiveness, is critical for the activation of innate immune function of lamellocytes.

## Discussion

In this study, we characterized the metabolic characteristics of *Drosophila* larval hemocytes and identified that these cells primarily rely on oxidative phosphorylation (OXPHOS) for energy metabolism. We showed that OXPHOS activity scales linearly with hemocyte proliferation during development and in response to genetic perturbations, except crystal cells. Furthermore, lamellocyte differentiation induced either by the hemocyte-specific expression of *hop^TumL^* or by parasitic wasp infestation significantly enhances both OXPHOS capacity and activity. This metabolic shift was accompanied by mitochondrial remodeling and the upregulation of trehalose metabolism-related genes. While trehalose metabolism was required for an effective melanization response, it did not significantly modify the differentiation of lamellocytes. Collectively, our findings indicate OXPHOS-dependent respiration, and its associated metabolic pathways are closely associated with hemocyte proliferation and innate immune activation, which imply a distinctive metabolic strategy of myeloid-like immune cells in *Drosophila*.

During larval development, hemocytes undergo substantial changes in mitochondrial metabolism that may reflect their distinct physiological requirements at different developmental stages. Upon hatching, approximately 200–700 hemocytes, predominantly plasmatocytes, are released into circulation and undergo extensive proliferation throughout larval development to reach 6,000–8,000 cells^14,22,55^. Notably, ATP production peaks at 96 h AEL, corresponding to a period of active larval feeding and hemocyte expansion. This increased metabolic activity may support energetic demands associated with hemocyte proliferation and maturation. Consistent with this possibility, mitochondrial number and size change between 72 and 96 h AEL, coinciding with the highest metabolic activity observed at 96 h AEL. However, despite a significant reduction in mitochondrial respiration and ATP production at 120 h AEL, mitochondrial number and size remain largely unchanged compared with those in 96 h AEL. This discrepancy suggests that mitochondrial morphology does not necessarily reflect metabolic activity and that mitochondrial function may be regulated independently of mitochondrial structure. Given that larvae gradually reduce feeding activity as they approach pupariation, the decline in hemocyte metabolism at 120 h AEL may reflect reduced nutrient uptake and systemic metabolism during late larval stages. Together, our observations suggest that hemocyte mitochondrial metabolism is dynamically regulated according to developmental and physiological requirements, and metabolic activity may represent related but distinct aspects of hemocyte development.

Macrophages serve as sentinels of cellular stress and immune challenge, exhibiting remarkable plasticity in response to various environmental stimuli^69^. In vertebrates, activated macrophages are traditionally categorized into pro-inflammatory M1 and anti-inflammatory M2 phenotypes, which display profound metabolic divergence. M1 macrophages preferentially utilize aerobic glycolysis and the pentose phosphate pathway (PPP) to support rapid energy production and the biosynthesis of inflammatory intermediates^70,71^, whereas M2 macrophages rely on OXPHOS^72^. In this context, our results demonstrate that *Drosophila* hemocytes, mainly consisting of macrophage-like plasmatocytes, show a metabolic signature similar to that of M2 macrophages, characterized by a high dependence on OXPHOS-driven metabolism. Interestingly, and in contrast to the vertebrate M1 paradigm, *Drosophila* hemocytes do not undergo a significant glycolytic shift even under immune challenge. Instead, they elevate total ATP production by the upregulation of OXPHOS capacity. By maintaining OXPHOS, hemocytes may be able to achieve the high metabolic efficiency required to support cellular remodeling of lamellocytes and support the synthesis of immune effectors for the melanization cascade. Alternatively, hemocytes may engage glycolysis transiently to mount a rapid response to acute infection and then revert to OXPHOS during the post-acute phase. This program may further support immune functions by directing carbon flux into the PPP pathway to maintain redox homeostasis^73^. Our data suggest the possibility that insect immune cells favor metabolic economy over the metabolic wastefulness of the Warburg effect. This strategy may be advantageous in the open hemolymph system where circulating sugar supports both systemic growth and immunity as a shared limiting source. Thus, immune activation may not universally require a glycolytic shift but may instead depend on the organism’s life traits and systemic metabolic preference.

Previous studies have shown that larval hemocytes are metabolically quiescent at steady-state, unchallenged conditions. However, sensory neurons triggered by wasp odors enhance GABA release to mount an immune response via lamellocyte differentiation^74^. The same pathway also modulates the ROS levels through succinate conversion^61^, which all contribute to metabolic activities. These studies indicate that TCA cycle may not be fully active in hemocytes under unchallenged conditions, which likely attenuates TCA cycle during the surveillance phase and constrains baseline metabolic activity of hemocytes. This strategy may be important for minimizing the systemic energy cost of maintaining a large immune cell population. Interestingly, this hypothesis is also consistent with studies demonstrating that a high sugar diet is detrimental to the development and immunity of larval hemocytes^55,75^. These collective findings, together with our analyses, support a hypothesis that larval hemocytes maintain a tightly regulated metabolic baseline that prioritize energy conservation during homeostasis while preserving sufficient bioenergetic plasticity for rapid metabolic reprogramming upon pathogen encounter.

In the late third instar, plasmatocytes comprise ∼ 95 % of hemocytes. Accordingly, our measurements in wildtype animals primarily reflect the metabolic dynamics of plasmatocyte population, with a minor contribution from crystal cells. Despite efforts to profile each hemocyte type in isolation, it was not feasible to completely separate the three lineages. Instead, we leveraged genetic backgrounds that bias expansion of specific hemocyte types, including *Hop^TumL^*, *Ras^v12^*, and *Notch^ICD^*, to enrich for the population of interest. Expression of *Ras^v12^* robustly expanded plasmatocytes by 10 to 50 folds with minimal lamellocyte expansion, whereas *Hop^TumL^* or wasp infestation enabled metabolic quantification of lamellocytes alongside activated plasmatocytes^16,24^. *Notch^ICD^* expression increased crystal cell differentiation but likely included a residual plasmatocyte fraction^13^. Although informative, these enrichment strategies are subject to admixture-related caveats because residual plasmatocytes may contribute substantially to the analyzed cells. Moreover, genetic manipulation of specific signaling pathways may induce immune activation in plasmatocytes that potentially influence the observed metabolic readouts. It is also worth noting that activation of genetically induced signaling pathways, including RAS-MAPK, JAK-STAT, or Notch pathway, may induce different metabolic adaptations despite producing similar metabolic readouts. Future cell-type resolved approaches, including advanced cell-sorting or single-cell metabolomic profiling, will be essential to define these programs and the underlying regulatory mechanisms across fully resolved hemocyte types and hematopoietic origins. Moreover, incorporating single-cell resolution analysis and space-and time-resolved measurements after immune or genetic perturbation will further clarify whether observed shifts reflect intrinsic metabolic rewiring or context-dependent activation.

While plasmatocytes in the unchallenged state express basal levels of carbohydrate transporters, lamellocytes significantly induced trehalose transporters and trehalase upon differentiation^5–7^. In addition, an elegant ^13^C-isotope labeling approach showed that activated hemocytes channel carbon sources primarily into the cyclic pentose phosphate pathway (cyclic PPP) to generate NADPH, together with ATP, pyruvate, or pentose^73^. Integrating the prior work with our analyses, we propose that activated plasmatocytes and lamellocytes maximize cyclic PPP flux to bolster antioxidant buffering for host protection, while sustaining viability via OXPHOS-mediated energy metabolism. By contrast, increasing the number of crystal cells did not significantly alter overall metabolic parameters. Although crystal cells were viable at the time of measurement, they may have adjusted their metabolic program in response to environmental changes *in vitro*. Moreover, *N^ICD^*-induced crystal cells may not fully recapitulate developmentally programmed crystal cell differentiation. Given that crystal cells are non-mitotic and expand by transdifferentiation^76^ and that crystal cells express a significant number of oxygen-carrying proteins^14^, these cells may adopt a distinct metabolic strategy relative to lamellocytes or plasmatocytes.

## Materials and methods

### *Drosophila* melanogaster stocks and genetics

The following *Drosophila* stocks were used in this study: *Hml*^Δ^*-Gal4* (S.Sinenko), *UAS-Ras^v12^* (BL4847), *UAS-hid, rpr* (Nambu J. R.), *UAS-hop^Tum-l^* (K.Brueckner), *Srp-Gal4* (L.Waltzer), *UAS-mito-HA-GFP* (BL8442), *Drp1 RNAi* (BL27682), *Hex-A RNAi* (BL35155), *Treh RNAi* (BL51810), *Pdk RNAi* (BL28635), *ND75 RNAi* (VDRC100733), and *UAS-Notch^ICD^* (U. Banerjee).

Fly stocks used in this study were maintained at 18 °C. Unless otherwise indicated, Gal4-UAS fly crosses were incubated at 25 °C and 70 % humidity. To synchronize larval development, approximately one hundred adult flies were allowed to lay eggs on a grape-juice agar plate for two-hour period. At 23 hours after egg laying (AEL), any hatched larvae were eliminated from the plate. Larvae were then transferred to and maintained on standard cornmeal-based food at 24 h AEL.

### Hemocyte isolation and immunohistochemistry

The following primary antibodies were used in this study: anti-Pxn (1:1,000, rabbit), anti-PPO2 (1:1000, rabbit), anti-pH3 (Sigma-Aldrich, 06-570, 1:500). Before hemocyte isolation, larvae were vortexed with glass beads (Sigma-Aldrich, G9268) for three minutes. This step was performed to extract total hemocytes, including sessile and circulating populations. Vortexed larvae were bled onto a slide glass (Immuno-Cell Int, 61.100.17) which was kept at 4 °C (or on ice) for 45 minutes to allow hemocytes to settle and adhere. Samples were fixed with 3.7 % formaldehyde in 1x PBS for 30 minutes. Following fixation, samples were washed three times in 0.4 % Triton X-100 in 1× PBS for 10 minutes and blocked in 10 % normal goat serum (NGS) in 0.4 % TritonX, 1× PBS for 30 minutes. The primary antibody was added, and samples were incubated in a humidified chamber overnight at 4 °C. The following day, samples were washed three times in 0.4 % Triton X in 1× PBS and incubated in the secondary antibody (Jakson Laboratory, 1:250) or Phalloidin (ThermoFisher, A12380, 1:400) for 3 hours at room temperature^77^. After washing 3 times with 0.4% Triton X in 1× PBS, samples were mounted using Vectashield (Vector Laboratory) with DAPI and imaged using the Zeiss Axio Imager M2 or Zeiss LSM900 with Airyscan confocal microscope. Images were analyzed by Image J (Version 1.54) or Imaris (Bitplane).

### Analysis of *D. melanogaster* samples using the Agilent Seahorse XFe96 analyzer

The Agilent Seahorse XFe96 analyzer was preset and maintained at 28 °C (heater on). The day prior to the analysis, the Agilent Seahorse XFe96 cartridge (Agilent, 102416-100) was hydrated using a utility plate (Agilent, 102416-100) filled with 200 μl of cell culture grade sterile water and incubated in a 28 °C non-CO_2_ chamber. Simultaneously, the Seahorse XF calibrant buffer (Agilent, 102416-100) was pre-warmed in 28 °C non-CO_2_ chamber. The following day, water was removed from both the utility plate and the cartridge. The utility plate was refilled with 200 μl of pre-warmed calibrant buffer in each well. The cartridge was combined with utility plate and incubated in the 28 °C non-CO_2_ chamber over 30 minutes before running the analyzer.

Larval brain was dissected in phosphate-buffered saline (PBS) and loaded onto the Agilent Seahorse XF96 cell culture microplates (101085-004) containing assay media. The brains were carefully settled to the bottom center of each well in the microplate. To ensure optimal signal, we recommend to gently put the brain to the center of the well between the three spherical supports using a pipet tip. For total hemocyte isolation, larvae were vortexed with glass beads (Sigma-Aldrich, G9268) for 3 minutes before bleeding. To isolate circulating and sessile hemocytes separately, larvae were first bled without vortexing to collect circulating hemocytes. The larval carcasses were then transferred to fresh Schneider’s *Drosophila* Medium (Thermo Fisher Scientific, 21720024) and gently scraped with forceps to release sessile hemocytes.

Hemocytes and S2 cells were prepared in Schneider’s *Drosophila* Medium and filtered using a 40 μm cell strainer (Falcon, 352340). Filtered cells were centrifuged at 6000 rpm for 5 minutes and suspended into the assay media. After adjusting the cell concentration to 1.5x10^6^ cells/ml, 180 μl of the assay media was added to each well.

The sample-loaded microplate was centrifuged at 3000 rpm for 3 minutes (Eppendorf, EP022628146) to ensure settling and adherence. The cartridge combined with utility plate was loaded with assay solution on injection ports and placed onto the Seahorse XFe96 analyzer and the run was initiated. The utility plate was replaced with the sample microplate after the equilibration phase.

### Assay media and solutions

Assay media were prepared immediately prior to each experiment based on the specific metabolic test. The base media for all assays were Seahorse XF DMEM medium, pH 7.4 (Agilent, 103575-100). For real-time ATP rate assay, glycolytic rate assay, and mito stress test, the base medium was supplemented with 10 mM of XF glucose (103577-100), 1mM of XF pyruvate (103578-100), and 2mM of XF glutamine (103579-100). For glycolysis stress test, the assay was performed in Seahorse XF DMEM medium, pH 7.4 mixed with 2mM of XF glutamine (103579-100).

Metabolic inhibitors or substrates were prepared as 10X stock solutions and diluted into the respective basal assay media: Oligomycin (Sigma-Aldrich, O4876, final concentration 1.5µM), 2-DG (Sigma-Aldrich, D8375, final concentration 50mM), Rotenone (Sigma-Aldrich, R8875 final concentration 0.5µM), Antimycin A (Sigma-Aldrich, A8674, final concentration 0.5µM) FCCP (Sigma-Aldrich, C2920 final concentration 10µM), glucose (Agilent Technologies, 103577-100, final concentration 10mM), or trehalose (Sigma-Aldrich, PHR1344, final concentration 5mM). These media were loaded into the Seahorse XFe96 analyzer injection ports immediately before the assay run.

### ATP assay

Circulating hemocytes were collected from larvae by bleeding into PBS and counted using a hemocytometer. ATP levels in hemocytes were measured using an ATP Determination Kit (Invitrogen, A22066), according to the manufacturer’s instructions. ATP levels were normalized to hemocyte number.

### Wasp infestation

*Drosophila melanogaster* larvae were collected at 72h AEL and exposed to *Leptopilina boulardi* (G486) wasps for 6 hours. Following the exposure, adult wasps were removed. The infestation process was maintained under controlled conditions of 25 °C and 70 % humidity. Infestation was confirmed by observation of wasp eggs or larvae inside the host, which was verified during bleeding. Encapsulation was assessed at 60 h post wasp infestation (hPI). Successful encapsulation was defined by the presence of a melanotic capsule. After scoring, each larva was dissected to confirm infestation. The encapsulation rate was calculated by the number of infested larvae with a melanotic capsule divided by the total number of infested larvae. The n represents the total number of larvae analyzed across all trials.

### TMRM staining

Circulating hemocytes from four larvae were collected by bleeding into 40μL of Schneider’s medium containing 100nM TMRM (Invitrogen, I34361) and incubated for 30 min at room temperature. Cells were washed once with 36μL of Schneider’s medium. After imaging, 4μL of 100μM FCCP diluted in Schneider’s medium was added to achieve a final concentration of 10μM and incubated for 5min before imaging again. TMRM fluorescence intensities were quantified within mitoGFP-positive pixels. For each image, the mean fluorescence intensity of a cell-free background region (F_0_) was subtracted from the measured TMRM fluorescence intensity.

### DHE staining

Larvae were bled and incubated with 30μM DHE in Schneider’s medium for 10min at room temperature. Hemocytes were then washed three times with Schneider’s medium for 3 min each. Next, samples were fixed in 3.7% formaldehyde in PBS for 3 min and washed once with PBS for 3 min. Finally, samples were mounted in VECTASHIELD (Vector Laboratory) with DAPI.

### Glycemic Level Assay

Glycemic levels, defined here as the combined circulating glucose and trehalose levels, were measured as previously described^78^, with modifications. Briefly, for each sample, 3 µL of hemolymph was collected from 15 larvae and diluted. An aliquot of the diluted hemolymph was reserved for protein quantification by Bradford assay, while the remaining sample was heated at 70 °C for 10 min to inactivate proteins. Samples were then centrifuged at 15,000 rpm for 5–10 min, and the resulting supernatant was collected and incubated with trehalase (Sigma-Aldrich, T8778) in trehalase buffer prepared as described by Tennessen et al. (2014) at 37 °C for 16–18 h. This treatment converts circulating trehalose to glucose; therefore, glucose measured following trehalase digestion represents the combined contribution of free glucose and trehalose and is reported here as the glycemic level. Following incubation, samples were centrifuged for 2 min, and total glucose levels in the supernatant were measured against glucose standards using a glucose assay kit (Sigma-Aldrich, GAHK20-1KT).

Where indicated, glycemic measurements were normalized to protein concentration determined from the reserved hemolymph aliquot using the Bradford assay (Bio-Rad, Cat. No. 5000006). Absorbance was measured using a FlexStation 3 Multi-Mode Microplate Reader. Data were processed using Microsoft Excel (Version 16.115) and plotted and analyzed using GraphPad Prism (Version 11.1.0).

### Succinate feeding

For succinate supplementation experiments, sodium succinate dibasic hexahydrate (Sigma, S2378) was added to dextrose-based fly food at a final concentration of 3% (w/v). Larvae were transferred from dextrose food to control or succinate-supplemented food 24 h before hemocyte collection.

### RT-qPCR

Hemocytes were collected by bleeding more than 50 larvae per sample. cDNA was synthesized using ReverTra Ace qPCR RT Master Mix (TOYOBO, FSQ-201). Quantitative PCR was performed using SYBR Green Realtime PCR Master Mix (TOYOBO, QPK-201) on a SteponePlus Real-Time PCR System (Thermo Fisher Scientific). Target gene expression was normalized to Rp49(Rpl32). Primer sequences are listed in Supplementary Table 1.

### Preprocess of scRNA-seq datasets

Drop-seq UMI count matrices of circulating hemocytes from 96 and 120 h AEL under steady state and wasp infection were retrieved from our previous study GSE141273 ^7^. Total of 12 libraries, three for each condition, were downloaded and merged. Because the gene annotation version was different between steady state (BDGP 6.22) and wasp infested (BDGP 6.02) data, an FBgn to Annotation ID conversion table from FlyBase (https://flybase.org; fbgn_annotation_ID_fb_2019_03.tsv) was used to convert gene IDs to the BDGP 6.22 version. Because these UMI matrices were already preprocessed (i.e., filtration of low-quality cells based on UMI, gene counts, and mitochondrial contents), no further filtration was performed. Briefly, a few outlier cells were first filtered using UMI thresholds; 96 and 120 h AEL wild type, UMI count > 50,000; 96 h AEL infested, > 70,000; 120 h AEL infested, > 55000. Then cells having UMI counts higher than two standard deviations from the mean UMI count were also removed to exclude possible multiplets. Low quality cells having only limited number of genes were also removed; 96 and 120 h AEL wild type, gene count < 200; 96 h AEL infested, < 300; 120 h AEL infested, < 400. Because mitochondrial contents in circulating hemocytes were a bit higher than those in lymph gland counterparts (a threshold of < 10% was used), we applied 20% as a threshold. Remaining cells were 1006 (96 h AEL wild type), 1671 (120 h AEL wild type), 5986 (96 h AEL infested), and 6693 cells (120 h AEL infested).

### Integration of scRNA-seq data from different conditions

For each dataset (96 h AEL steady state, 96 h AEL wasp infested, etc.), libraries were first predicted for cell types using a lymph gland from 72 to 120 h AEL as a reference as we performed in the previous study ^7^. In details, each dataset was first transformed to a Seurat object and three independent sequencing libraries were integrated using FindIntegrationAnchors() and IntegrateData() with default parameters ^79^. Then, cell type annotation was transferred from the lymph gland dataset using FindTransferAnchors() and TransferData() with default parameters. Non-hematopoietic cell types (dorsal vessel, neurons, ring gland, and posterior signaling center cells) and a few minor subclusters (< 0.1% of total population) were removed. In the end, 995 (96 h AEL steady state), 1356 (120 h AEL steady state), 5674 (96 h AEL wasp infested), 6376 cells (120 h AEL wasp infested) remained. After the prediction step, UMI counts were normalized, log-transformed, and scaled, and PCA analysis was performed which identified 40 significant principal components (PCs). Four datasets were integrated using Harmony ^80^ with default parameters. t-SNE and UMAP plots were generated using the selected numbers of PCs. Genes identified through the single-cell RNA sequencing analysis were further validated by quantitative real-time PCR (Supplementary Figure 8).

## Code availability

In-house Python and R source codes used in this study are publicly available on GitHub repository (https://github.com/sangho1130/dmel_metabolism). Analyses using public tools were performed using default parameters, otherwise, described in the Method section. All analyses were performed using Python (version 2.7.5), R (version 3.5.3), and R Studio (version 1.1.383).

## Data availability

All data generated during and/or analyzed during the current study are included in this published article and its supplementary files.

## Quantification of samples, statistics and reproducibility

All hemocyte samples were imaged using a Zeiss Axiocam 503 camera or a Zeiss LSM 900 confocal microscope. Hemocyte populations were quantified using ImageJ (version 1.54). Statistical analyses were performed using GraphPad Prism 11 (version 11.1.0). Bars in all graphs indicate means, and error bars represent standard deviation (SD) or standard error of the mean (SEM), as specified in the figure legends. Statistical significance was assessed using unpaired t-test. Significance levels are indicated as follows: n.s: not significant (p > 0.05). *p < 0.05; **p < 0.01; ***p < 0.001. All experiments were repeated at least twice.

## Acknowledgements

We thank all members of the Shim laboratory for their helpful comments. We thank Bloomington Drosophila Stock Center (NIH P40OD018537) and the Drosophila Genomics Resource Center (NIH 2P40OD010949), FlyBase (release 2025_04, 2019_03), and the Korean Drosophila Resource Center for supporting the fly community. This work was supported by R35GM128752-NIH NIGMS Kuni Foundation Discovery Grants for Cancer Research to Y.V.K. and Basic Science Research Program through the National Research Foundation of Korea (NRF) funded by the Ministry of Education (RS-2024-00349703, RS-2025-02232977, and RS-2026-25501101) and the New Faculty Startup Fund from Seoul National University to J.S.

## Author contributions

DL performed Seahorse analyses, wasp infestation analyses, hemocyte counting, imaging, and quantitative analyses. FK and NC performed wasp infestation analyses and associated hemocyte counting. SHY, KTL, and JWN performed *in silico* analysis using hemocyte single cell RNA sequencing. DL, YVK, and JS designed this study and wrote the manuscript. YVK and JS supervised the project and conceived the idea.

