## Supplementary Figures 1-8 for "Mitochondrial remodeling controls innate immune effector function in *Drosophila* hemocytes"

### Supplementary Figure 1

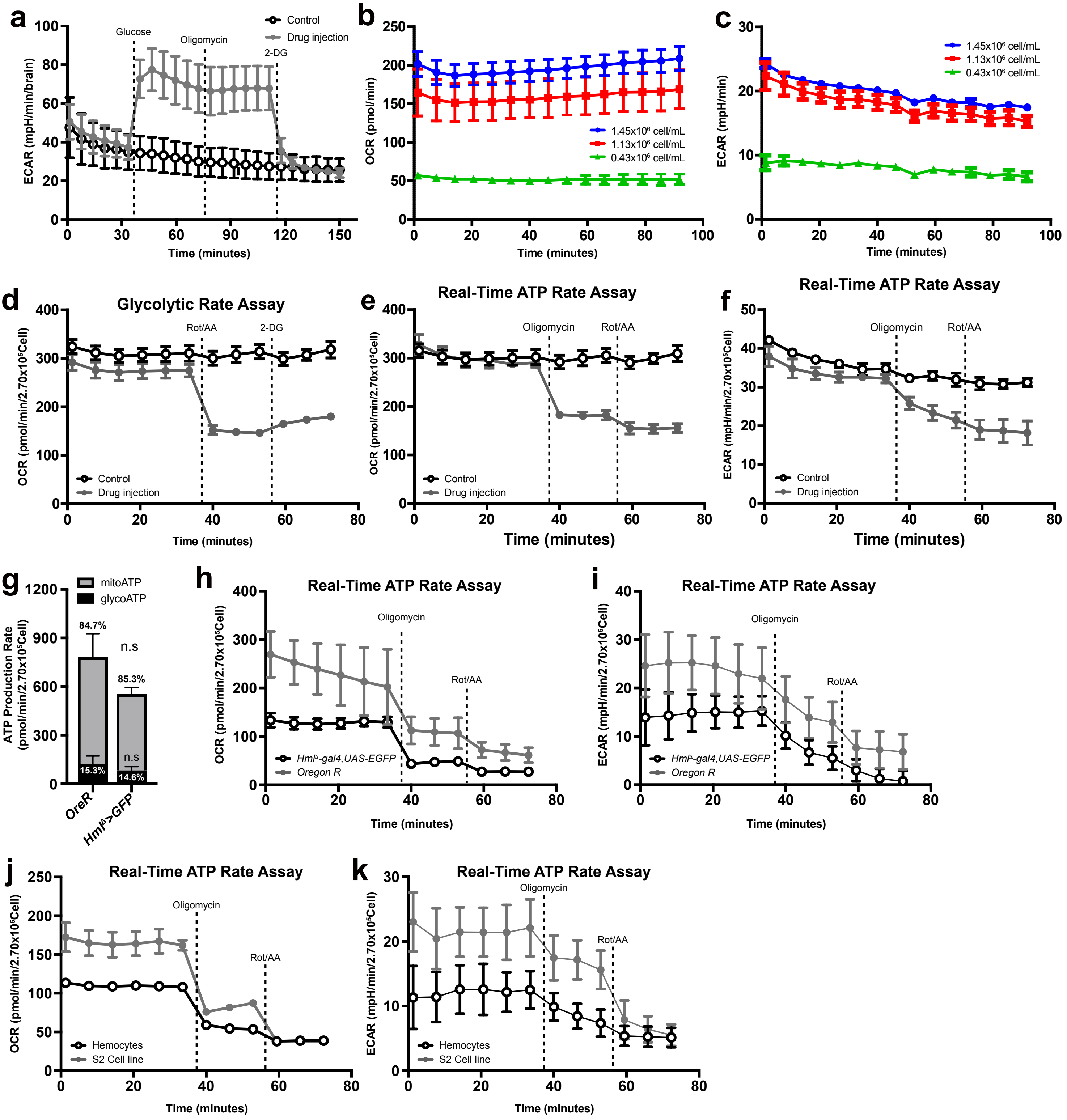

#### Supplementary Figure 3

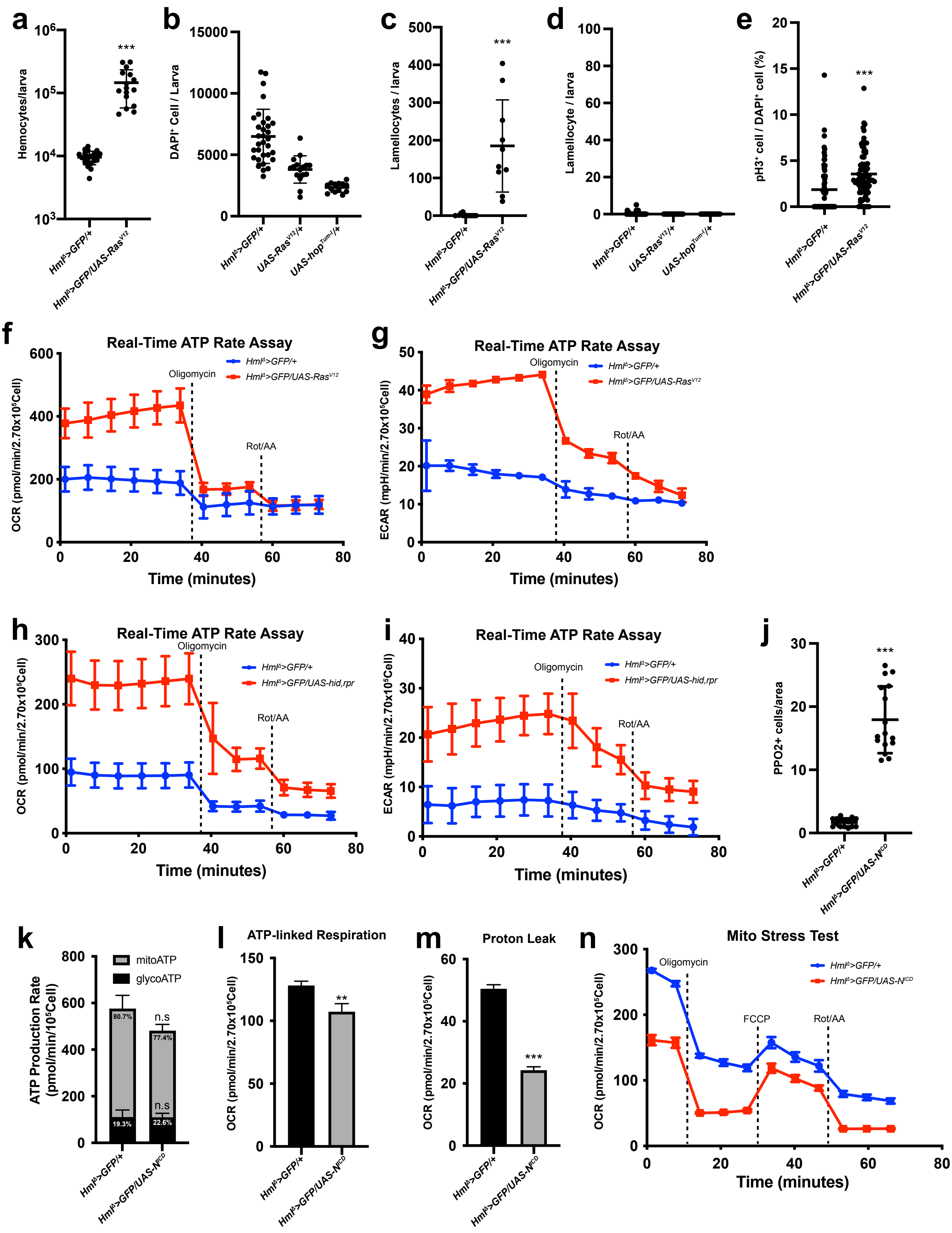

### Supplementary Figure 4

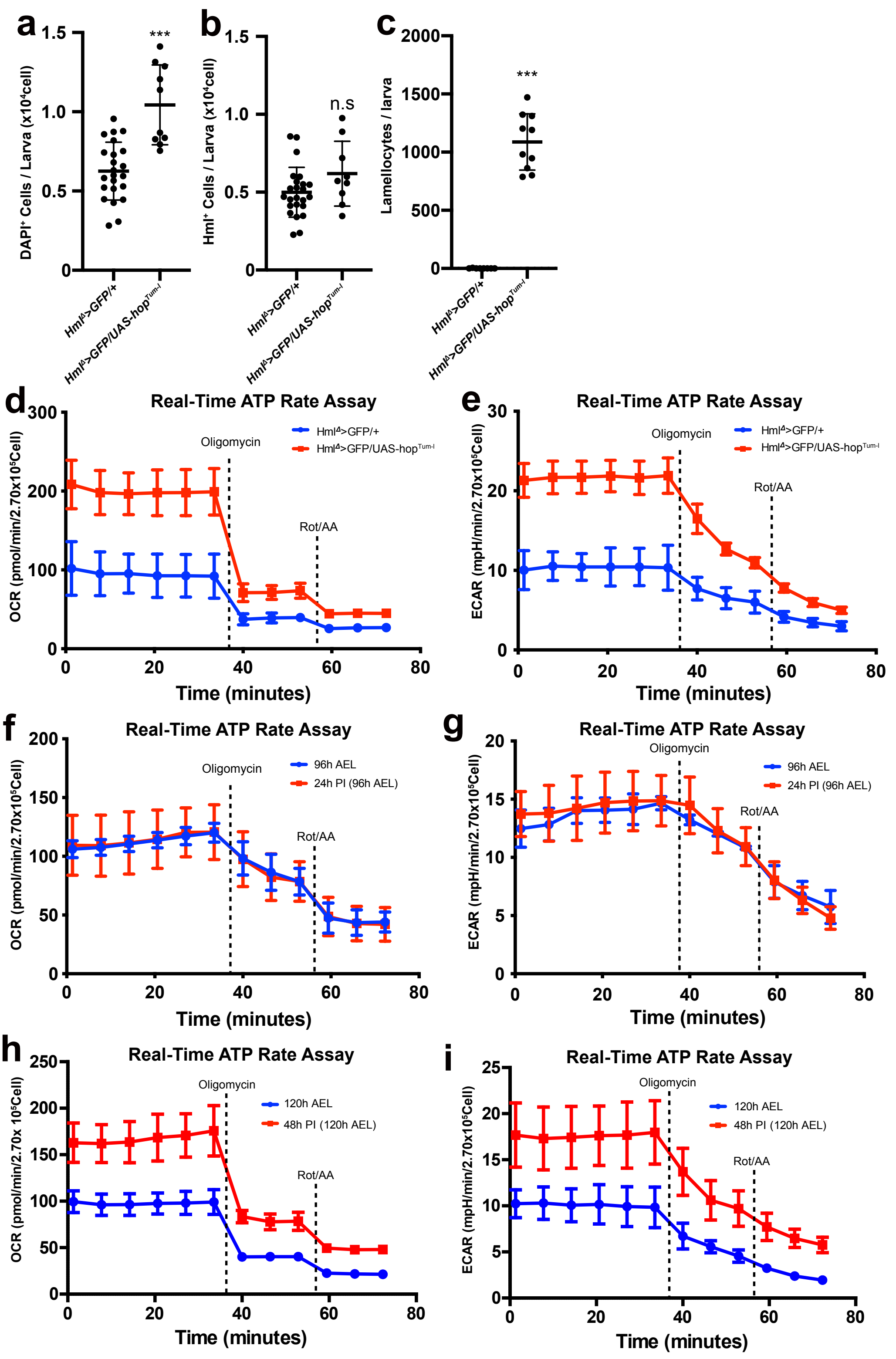

### Supplementary Figure 5

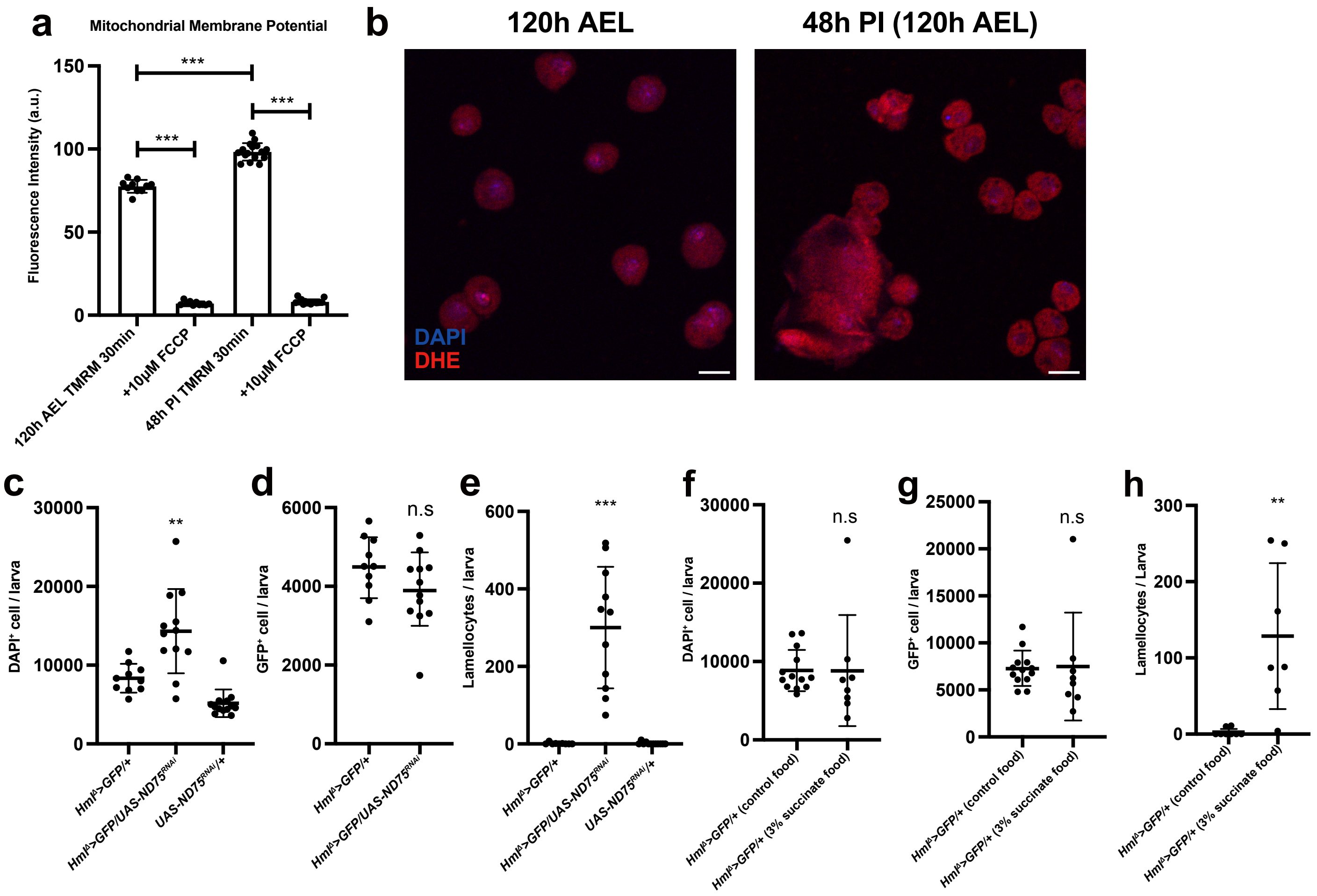

### Supplementary Figure 7

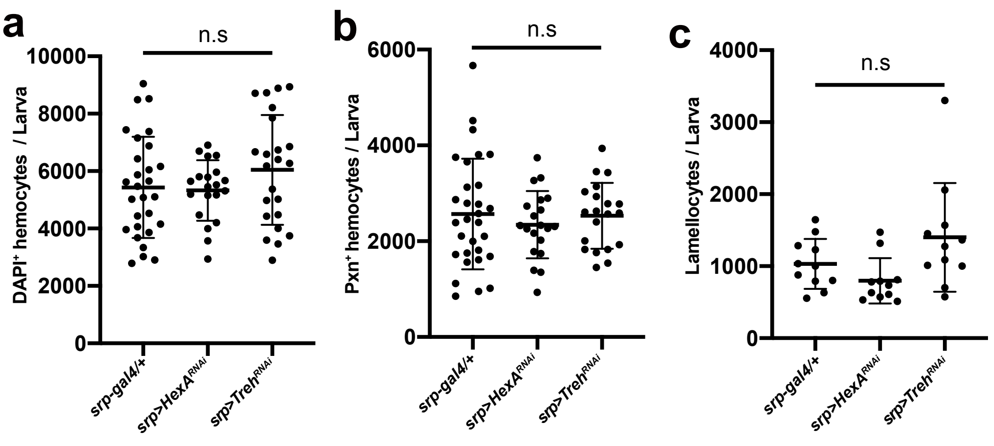

#### Supplementary Figure 8

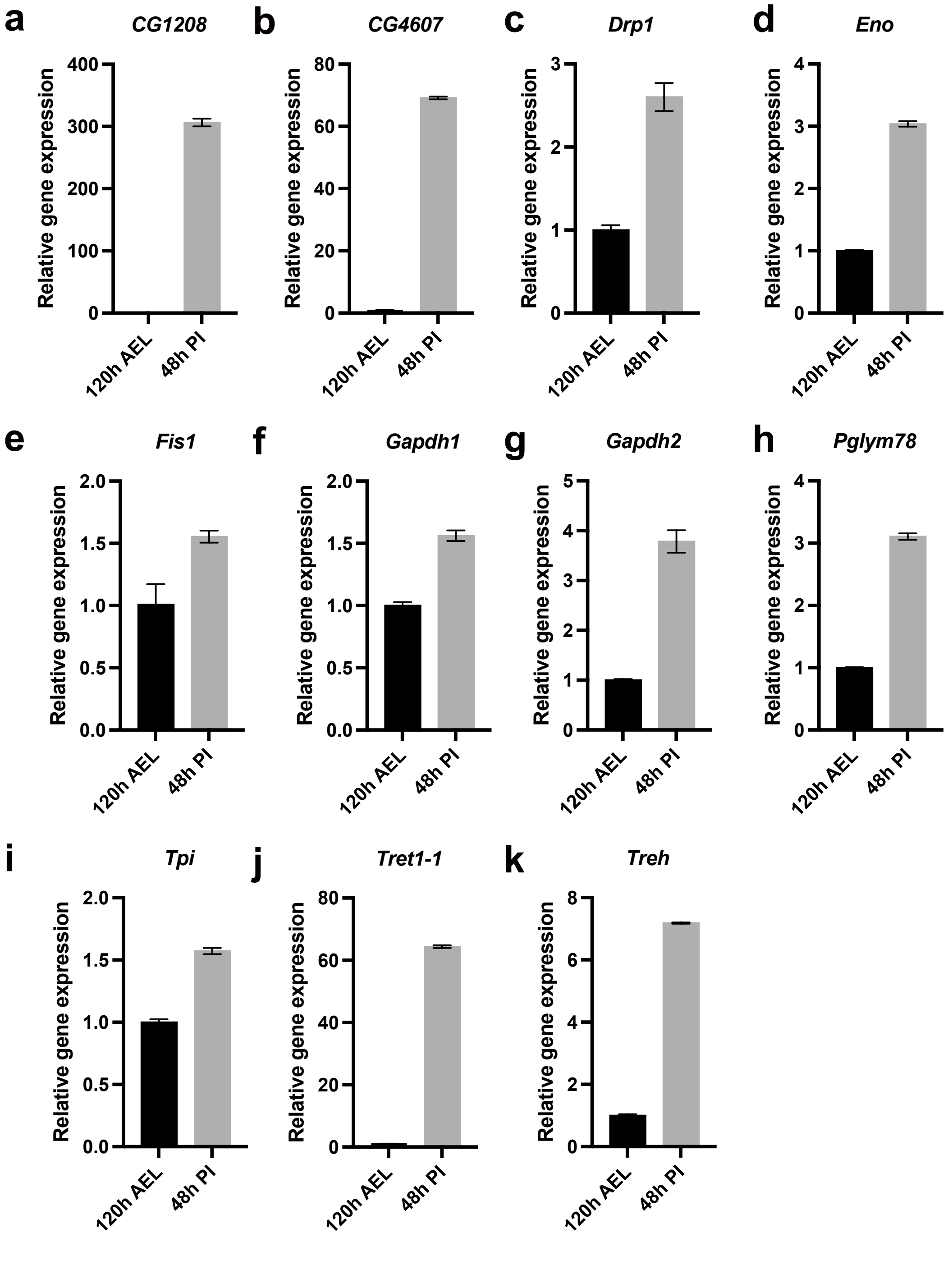
